# SEC-seq: Association of molecular signatures with antibody secretion in thousands of single human plasma cells

**DOI:** 10.1101/2022.08.25.505190

**Authors:** Rene Yu-Hong Cheng, Joseph de Rutte, Andee R. Ott, Lucienne Bosler, Wei-Ying Kuo, Jesse Liang, Brian E Hall, David J Rawlings, Dino Di Carlo, Richard G. James

**Author notes:** Contributed equally to this study (Rene Yu-Hong Cheng, Joseph de Rutte).

## Abstract

Protein secretion drives many functions *in vivo*; however, methods to link secretions with surface markers and transcriptomes have been lacking. By accumulating secretions close to secreting cells held within cavity-containing hydrogel nanovials, we demonstrate workflows to analyze the amount of IgG secreted from single human antibody-secreting cells and link this information to surface marker expression and transcriptional profiles from the same cells. Measurements using flow cytometry and imaging flow cytometry corroborated an association between levels of IgG secretion and CD138 expression. Using oligonucleotide-labeled antibodies and droplet-based sequencing, we show that pathways encoding protein localization to the endoplasmic reticulum, NADH complex assembly, and mitochondrial respiration were most associated with high IgG secretion. Altogether, this method links secretion information to cell surface and single-cell sequencing information (SEC-seq) and enables exploration of links between genome and secretory function, laying the foundation for numerous discoveries in immunology, stem cell biology, and beyond.

## INTRODUCTION

Organisms critically depend on the proteins or other factors which cells secrete into their environment that can act locally in a paracrine manner or systemically. For example, one of the main roles of B cells is to respond to antigens with the production and secretion of large quantities of immunoglobulins targeting antigen epitopes. In this process, antibody-secreting B cells differentiate, undergoing significant phenotypic, morphological, and genetic changes. Linking these functional changes in the capacity for secretion of immunoglobulins to genetic/phenotypic profiles at the single-cell level can uncover the population heterogeneity and potential new cell states.

Standard single-cell analysis tools cannot simultaneously quantify external cellular information (e.g., secreted proteins) with cell surface and/or intracellular information. One possible method to infer information about secretions is to analyze the intraceullular concentration of the transcripts (single cell transcriptomes) or proteins (flow cytometry) that potentially will be secreted. While transcripts associated with secreted proteins may sometimes correlate to secretion^1–3^, transcript levels cannot adequately account for several post-transcriptional activities such as RNA splicing, translation, post-translational modifications, enzymatic cleavage, or even storage of secreted proteins in secretory vesicles prior to secretion. Furthermore, intracellular proteins, including secreted proteins, can be analyzed, e.g. using intracellular cytokine staining, but through a destructive process that involves permeabilization and fixation of the cell. Forming pores in the cell is necessary for fluorescently-labeled antibodies to penetrate the intracellular space and bind to chemically fixed proteins. Downsides of permeabilization and fixation include the loss of cell viability and destruction of other intracellular molecules, such as mRNA, which limits downstream transcriptomic analysis. Finally, the presence of secreted proteins on or within the cell does not necessarily indicate that these proteins would have been secreted.

Other tools to characterize cell-secreted products lack the quantitative resolution, throughput, and multiplexing of flow cytometry and do not directly link secretions to transcriptomic information. Researchers have utilized optofluidic pens or other microfluidic compartments to isolate single cells and accumulate secreted products for analysis on solid surfaces near the cells. Two recent instruments that employ these approaches are the Beacon system from Berkeley Lights, and Isoplexis’ Isolight system. Both systems use microscopic imaging to analyze the secreted products from cells, with dynamic range and the number of color channels constrained by the cameras and filter sets used^4–6^. In contrast, the gold standard in single-cell analysis, flow cytometers, leverage laser-based excitation and PMT-based detection to achieve sensitive multiplexed measurements with high dynamic range. Surface markers are not readily accessible in the Isolight system, while the Beacon system can analyze a few. The number of cells that can be analyzed depends on the number of optofluidic pens or microchambers on a single chip, usually between 1,000 to 10,000. While the Beacon system can sort cells after analysis, this is not achievable with the Isolight system. Another assay format that is compatible with flow cytometry uses the cell’s surface itself to capture secreted products (e.g. the cytokine-catch assay from Miltenyi). This assay format requires specialized bi-specific antibodies that bind to CD45 on leukocytes and capture specific cytokines but is not easily extensible to other secreted products or cells that lack consistent or high levels of CD45, including plasma B cells. Secretions can also diffuse away and bind to neighboring cells, leading to crosstalk since cells are not confined in individual compartments^7^. No current technology has been able to link the amount and type of secreted molecules from a single cell with its transcriptional profile, nor has this type of association been multiplexed at a scale of hundreds to thousands of single cells simultaneously.

We demonstrate a technology that overcomes these tradeoffs, combining the multiplexing and high-throughput quantitative analysis of surface markers by flow cytometry or transcripts by single-cell RNA-seq with the ability to quantify secretions (IgG) in the same single cells. The approach uses microscale hydrogel particles with a bowl-shaped cavity, called nanovials^8^, which capture cells and their secretions and are compatible with flow cytometry and single-cell sequencing instruments. To study secretions of antibody-secreting cells, we apply this technology to achieve an 8-plex multiplexed secretion assay, including 6 channels dedicated to cell surface markers, 1 channel to cell viability, and a final channel to IgG secretion. The approach was compatible with fluorescence activated cell sorting (FACS) using the Nanocellect WOLF and imaging flow cytometry using the Amnis ImageStream^®X^. By using oligonucleotide-barcoded antibodies, we could also link IgG secretion directly to transcriptomes in the same cells by introducing nanovials containing antibody secreting cells directly through a 10X Chromium single-cell RNA-seq workflow. This new method, SEC-seq has enabled us to learn several things about the biology of antibody-secreting B cells and showed that transcripts involved in antibody production/metabolism rather than the antibody itself, were most highly associated with high secretion. We envision that this method could help us better understand the determinants of protein secretion in human antibody-secreting cells and other models, and better engineer secretory function, unlocking new therapeutic modalities.

## RESULTS

### Workflow for quantifying IgG secretion by human plasma cells

We first developed a workflow for loading and analyzing the secretions of single human B cells adhered to hydrogel nanovial particles (Fig. 1). This workflow involves capturing single cells into nanovial cavities by linking nanovials to conjugated antibodies targeted to surface proteins. Once captured into nanovials, cells are incubated to facilitate the accumulation of secreted IgG onto anti-IgG antibodies pre-coated to the cell-associated nanovial surface. The nanovial-bound secreted IgG and other cell surface markers are then tagged with antibodies with either fluorescent labels or oligonucleotide barcodes. Labeled nanovials and cells are then analyzed using flow cytometry, imaging flow cytometry, or sorted by FACS and analyzed by single-cell sequencing.

**Figure 1.**
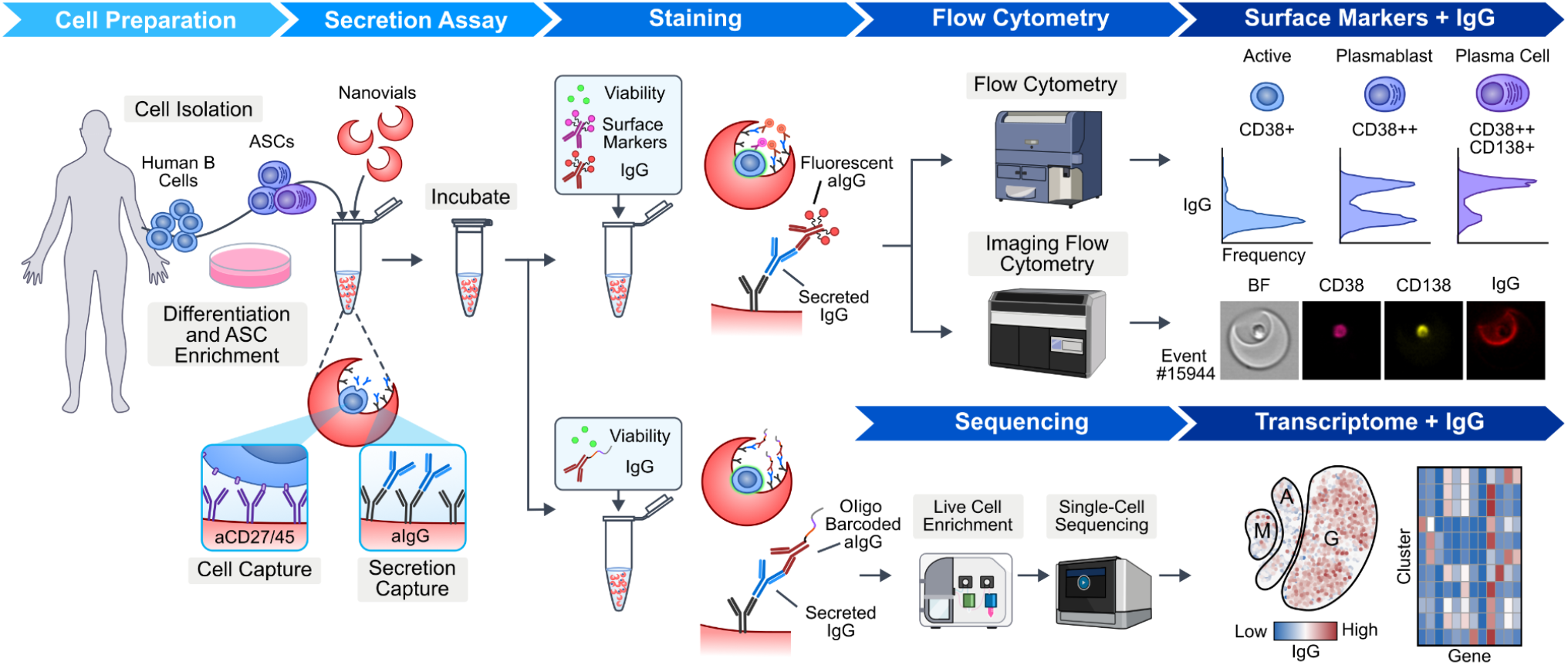
Workflow to link IgG secretion to surface markers and transcriptomes at the single-cell level. Human B cells are isolated from donors and expanded in a differentiation cocktail to enrich antibody secreting cells (ASCs). Cells are then loaded into a slurry of nanovials in a tube where they bind to antibodies on the nanovials against cell surface markers (CD27, CD45). Cells are incubated to accumulate secreted IgG on the nanovial surface via anti-IgG capture antibodies, and then nanovials are stained with fluorescent or oligo-barcoded anti-IgG. Cells associated with the nanovials are also stained with viability and other surface marker stains. Stained nanovials and associated cells are analyzed by flow cytometry (LSR II flow cytometer) or imaging flow cytometry (ImageStream) or sorted (Nanocellect WOLF) for single-cell transcriptomics using the 10X Chromium system. Data linking IgG secretion with surface markers and transcriptomes at the single-cell level is acquired and analyzed.

To identify optimal surface markers to capture *ex vivo* differentiated human plasmablast/plasma cells^9^ into nanovials, we tested a panel of antibodies against surface proteins (CD45, CD27, CD38, CD31) expressed in B cells and analyzed capture by flow cytometry. Nanovials are made of highly transparent hydrogel, however, the shape and larger size leads to a unique scatter signature that is readily distinguished from other cell events^10^. Cells loaded on nanovials were discriminated from free cells and empty nanovials using a combination of flow cytometry scatter and fluorescence gating. Based on live-dead stain, only live cells were gated in the downstream analysis (Fig. S1a). We found that antibodies against CD27 yielded the highest percentage of total cells captured, and the captured cells represented a broad range of cell types, including CD19^high^ active B cells, as well as CD19^low^ IgM^+^ cells, IgA^+^ cells, and IgM/IgA double negative cells (Fig. S1b).

### Quantifying IgG secretion in plasma cells expressing different cell surface markers

Previously, we found heterogeneity in antibody secretion rate for plasma cells measured using ELISPOT (Fig. S2), however, we were unable to investigate these sub-populations further due to the inability to characterize other properties of cells in the ELISPOT format. Using nanovials, we can use flow cytometry to simultaneously quantify IgG secretion along with cell surface and intracellular proteins at the single-cell level. To explore the heterogeneity of phenotype in human antibody-secreting cells, we isolated B cells from peripheral blood mononuclear cells (PBMCs), and differentiated these into heterogeneous populations of plasma cells, plasmablasts and activated B cells *ex vivo*^*9*^ (see methods for more details). After differentiation, we loaded cells onto 55 µm-diameter nanovials functionalized with anti-CD27 and anti-human IgG, and stained cells with a panel of B cell/plasma cell and immunoglobulin class surface markers to define sub-populations, as well as anti-IgG for detecting secreted IgG on nanovials (Fig. 2a). A large fraction of loaded B cells exhibited a phenotype consistent with plasmablasts (PBs) or plasma cells (PCs; CD19^lo^CD38^+^), which may represent antibody secreting cells (ASCs). A small portion of loaded cells exhibited a phenotype of activated B cells (CD19^hi^CD38^lo^). The ASCs could further be categorized by antibody isotype. We observed sub-populations of cells with surface expression of IgM or IgA, and double negative (DN) cells that are most likely IgG^+^ ASCs; few IgE cells are present in our culture system (Fig. 2b).

**Figure 2.**
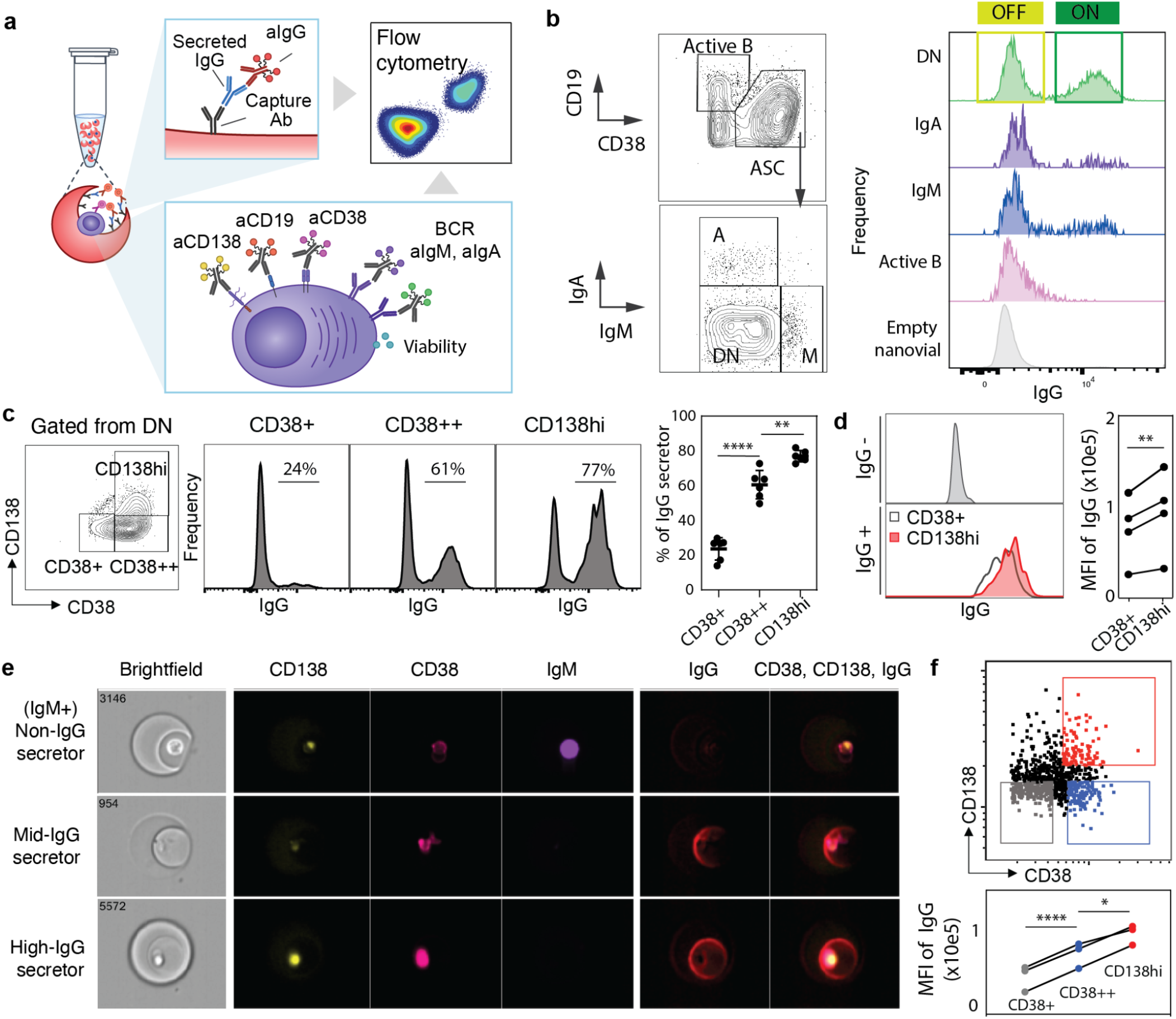
Linking IgG secretion to cell surface markers and intracellular machinery using flow cytometry. (**a**) Schematic of staining format used to analyze single B cell IgG secretion and cell surface markers using flow cytometry. (**b**) Flow cytometry density scatter plots for surface markers to identify populations of ASCs from active B cells (n=999) using CD19 and CD38 staining. IgA cells (n=319), IgM cells (n=436), and ASCs not producing either IgA or IgM (double negative, DN, n=6004)) were gated based on IgA and IgM staining. Fluorescence histograms of IgG secretion signal for the various identified gates and empty nanovials containing no cells. (**c**) Contour plot for CD38 and CD138 staining in the DN population and fluorescence histograms of IgG signal from populations within these gates. Dot graph presents the % of IgG secretors from each ASCs subset (n = 5, from 2 donors). To assess significance, we used a paired one-way ANOVA with Tukey’s multiple comparison test (* p < 0.05, ** p <0.01, *** p < 0.001). (**d**) Histogram of IgG levels for IgG non-secretors and IgG secretors from different ASC subsets. Linked dot graph presents the mean fluorescence intensity (MFI) of IgG secretors from each ASC subset. A two-sided paired t-test is performed for each experiment (n = 4, from 1 donor); significance is under a significance level of α = 0.05. (* p < 0.05, ** p <0.01, *** p < 0.001). (**e**) Imaging cytometry results confirm that IgM^+^ cells have low levels of secreted IgG, CD138low and CD38low populations have intermediate levels of secreted IgG, and CD138^+^CD38^++^ have the highest levels of secreted IgG. (**f**) Imaging flow cytometry gating strategy for IgG quantification by cell subtype (upper). Dot graph presents the MFI of IgG from each ASC subset (n = 3, from 3 individual donors). To assess significance, we used a paired one-way ANOVA with Tukey’s multiple comparison test (* p < 0.05, ** p <0.01, *** p < 0.001).

We next associated IgG secretion with the different B cell subtypes. As expected, activated B cells and the majority of IgA and IgM cells exhibited little to no IgG secretion (Fig. 2b). To address the potential cross-reactivity and diminish the signal from free cells in the loading step, we included unbound anti-IgG antibody in solution during the cell loading procedure to block these interactions. After making these changes to the protocol, we showed that IgM^+^ cells had reduced IgG^+^ signal (Fig. S3). A large percentage of DN cells exhibited high levels of IgG secretion (Fig. 2b-c). However, the distribution of IgG secretion in DN cells was bi-modal, indicating that a large percentage of these cells did not produce IgG despite the fact that most expressed surface markers that are conventionally associated with ASCs (Fig. 2c).

To further investigate the surface markers associated with IgG secretors, we further gated DN cells based on thresholds of CD38 and CD138 (boxes, Fig. 2c), markers which increase in expression during PC maturation. Of the DN cells, we observed increased proportions of IgG secreting cells depending on the expression of PB/PC maturation markers: low in immature PBs (CD38^+^CD138^lo^; ∼20% IgG), intermediate in PBs (CD38^++^CD138^lo^; ∼60% IgG) and high in PCs (CD38^++^CD138^hi^; ∼80% IgG, Fig. 2c). Overall, these data showed that CD138 positivity was the best marker to identify a B cell as an IgG secretor or not (Fig. 2c). When we compared the mean fluorescence intensity (MFI) of IgG between PCs and immature PBs, we observed significant increases in the PCs (Fig. 2d). These data demonstrate that while PC phenotype corresponds to a large increase in the proportion of IgG secretors, there is also a small increase in secretion amount by PCs relative to immature PBs.

We used imaging flow cytometry (Amnis ImageStream) to confirm that the anti-IgG signals we observed in PCs originated from single cells and were secreted into the associated nanovial. We analyzed loaded B cells on ImageStream (Gating strategy, Fig. S4) and used the images to measure fluorescence on the nanovials and cells separately. The signal for secreted IgG was distinct from fluorescence in the cell and evident as a crescent/ring shape on the inner surface of nanovials (Fig. 2e). Upon quantification of IgG in the other cell phenotypes, we found that CD38^++^CD138^hi^ PCs exhibited significantly higher IgG^+^ signal relative to CD38^++^CD138^lo^ or CD38^+^ cells (Fig. 2f and representative images Fig. S5). Collectively, these data indicate that CD138^hi^ PCs are the predominant source of secreted IgG in human plasma cell cultures.

### Compatibility of nanovial analysis with single-cell sequencing

We first evaluated the general compatibility of nanovials with single-cell transcriptomic sequencing using the 10X Genomics Chromium system. The Chromium system uses a microfluidic droplet generator to encapsulate cells with single barcoded hydrogel beads inside drops, where lysis and single-cell reverse transcription is performed. The gel beads contain oligonucleotides that hybridize to mRNA and other feature barcodes from the cell sample that also comprise a unique barcode that can be associated with each single cell. In our modified workflow, we simply replaced the cell sample in the microfluidic chip with nanovials loaded with cells (Fig. 3a). Nanovials with a diameter of 35 µm could be introduced into the microfluidic chips and entered into microfluidically generated droplets (Fig. S6a-b). Nanovials of larger sizes could flow through the chips, but only after deforming significantly to pass through the channels (Fig. S6c), which led to clogging in some experiments. To reduce the chance of clogging, we used 35 µm-diameter nanovials for the remaining single-cell sequencing experiments.

**Figure 3.**
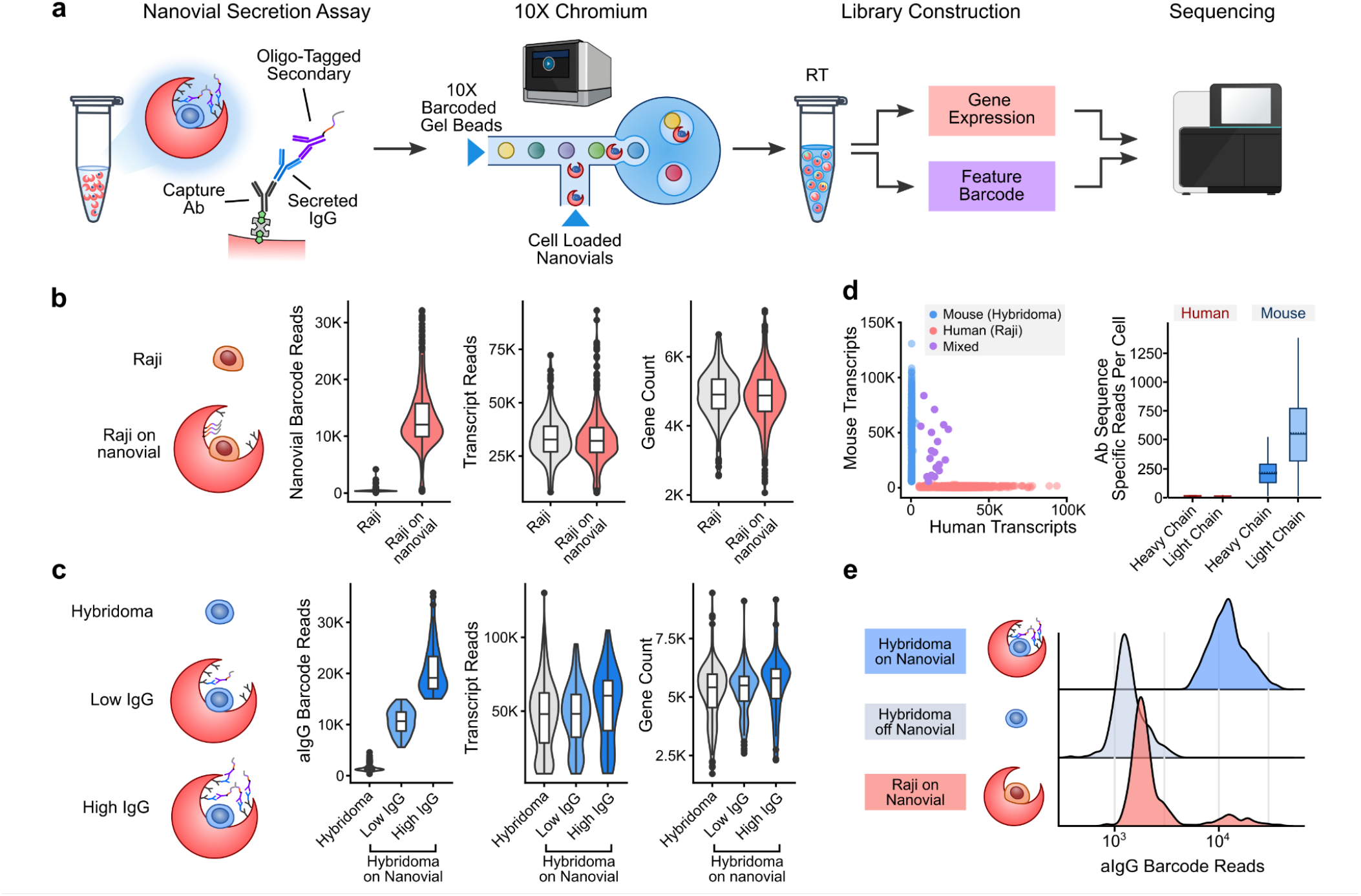
Compatibility of nanovials with single-cell transcriptomic sequencing using the 10X Genomics Chromium system. (**a**) Workflow for barcoding and analyzing secretions on nanovials. Captured secreted IgG is labeled with oligonucleotide-modified anti-IgG antibodies. Nanovials and associated cells are introduced into the 10X Chromium Next GEM Chip and emulsions are formed containing nanovials and Barcoded Gel Beads. Reverse transcription (RT) is performed to create cDNA and then amplified to form separate feature barcode and gene expression libraries. These libraries are sequenced, and feature barcode reads are linked to each cell’s transcriptomes. (**b**) Human Raji cells on nanovials (transcripts linked with a streptavidin feature barcode, Nanovial Barcode) are compared with Raji cells off nanovials based on the number of transcript reads and gene count. (**c**) Mouse hybridoma cells that secrete IgG that are off nanovials are compared to cells on nanovials based on the number of anti-IgG feature barcode reads, transcript reads, and gene count. (**d**) Scatter plot of mouse vs. human transcript counts when equal amounts of human Raji and mouse hybridoma cells on nanovials are input into the Chromium system. Specific heavy and light chain antibody sequences are recovered from mouse hybridoma cells. (**e**) Secreted IgG feature barcode read histograms for hybridomas on nanovials, hybridomas off nanovials, and Raji cells on nanovials.

We found that mRNA from mouse hybridoma cells and human Raji cells seeded on nanovials was successfully transcribed and sequenced. The number of transcripts and transcribed genes recovered from Raji cells on nanovials was comparable to freely suspended cells (Fig. 3b). Notably, nanovials can be tagged with feature barcodes (a nanovial barcode) by coating with oligonucleotide-conjugated streptavidin. We used this feature to differentiate Raji cells loaded on nanovials from cells freely floating in suspension (Fig. 3b). Similarly, we used reads of anti-IgG associated feature barcodes to categorize hybridoma cells as off nanovial, on nanovial with low anti-IgG barcode reads, or on nanovial with high anti-IgG barcode reads (Fig. 3c). Similar to our results with Raji cells, we found a comparable number of transcripts and gene numbers recovered from hybridoma cells on nanovials when compared to free hybridoma cells, which were largely independent of the amount of IgG secretion (Fig. 3c).

We next evaluated mixing by loading mouse hybridoma cells and human Raji cells on separate batches of nanovials and mixing the batches together before loading them into the microfluidic device. We prepared nanovial samples at the recommended concentration to recover 1000 - 2000 cells (10,000 nanovials assuming 10-20% cell loading). At this concentration we found the majority of cells had single species-specific gene reads (Fig. 3d), while a small minority (1.5%) had both human and mouse reads, suggesting coincident loading of two cell types in a droplet (Fig. S7). We searched the cDNA library for heavy and light chain sequences for the anti-hen egg lysozyme (HEL) antibody produced by the hybridomas. Sequences were recovered from the hybridoma population and were not detected in the Raji cells, as expected (Fig. 3d). Consequently, using nanovials did not lead to increased cell-cell mixing (i.e. shared barcodes for more than one cell) compared to statistical expectations.

By adding an oligonucleotide-barcoded anti-IgG label, we could link the secretion of IgG to the mouse transcriptome for individual cells on nanovials, an approach we refer to as secretion encoded cell sequencing (SEC-seq). As expected, the lowest number of anti-IgG feature barcode reads were associated with free cells (not loaded in nanovials) (Fig. 3e). In contrast, events with the highest IgG feature barcode reads were associated with mouse hybridomas on nanovials. As expected, most Raji cells on nanovials had low anti-IgG feature barcode reads, with a small group with higher reads proportional to the fraction of nanovial multiplet events (Fig. 3e, Fig. S7).

### Simultaneous quantification of protein secretion and single-cell transcriptome sequencing (SEC-seq)

After confirming the ability to perform linked secretion analysis and single-cell sequencing, we applied the SEC-seq technique to explore the transcriptomes of ex vivo differentiated human ASCs as a function of their IgG secretion phenotypes. In this workflow, we adapted the SEC-seq protocol by pre-sorting nanovials containing viable human B cells immediately prior to loading into emulsions with the 10X Barcoded Beads (Fig. 4a, and gating in Fig. S8a, Cell Ranger QC summary in Fig. S8b). The data from the sequencing was analyzed to simultaneously quantify IgG secretion (SEC) via signal from barcoded IgG antibodies (left panel, Fig. 4b) and gene expression sequencing (right panel, Fig. 4b). Clustering of the transcriptional data was largely driven by expression of the specific antibody isotypes (Fig. S9). The majority of IgG-secreting cells were in clusters expressing IGHG1, IGHG2, IGHG3 and IGHG4 (Fig. S9), with a small minority in clusters expressing IGHM, IGHA1. We did *not* observe a strong correlation between the quantity of UMIs and IgG secretion signals (Fig. 4c). Finally, factors conventionally associated with PB/PC differentiation (XBP1, IRF4, PRDM1) were expressed uniformly in all subclusters, regardless of antibody isotype identity or IgG secretion level (Fig. 4d). Because the strength of the secretion signal did not correlate with overall transcription or with levels of conventional PB/PC transcriptional activators, we reasoned that additional differentially expressed genes specifically regulate IgG secretion.

**Figure 4.**
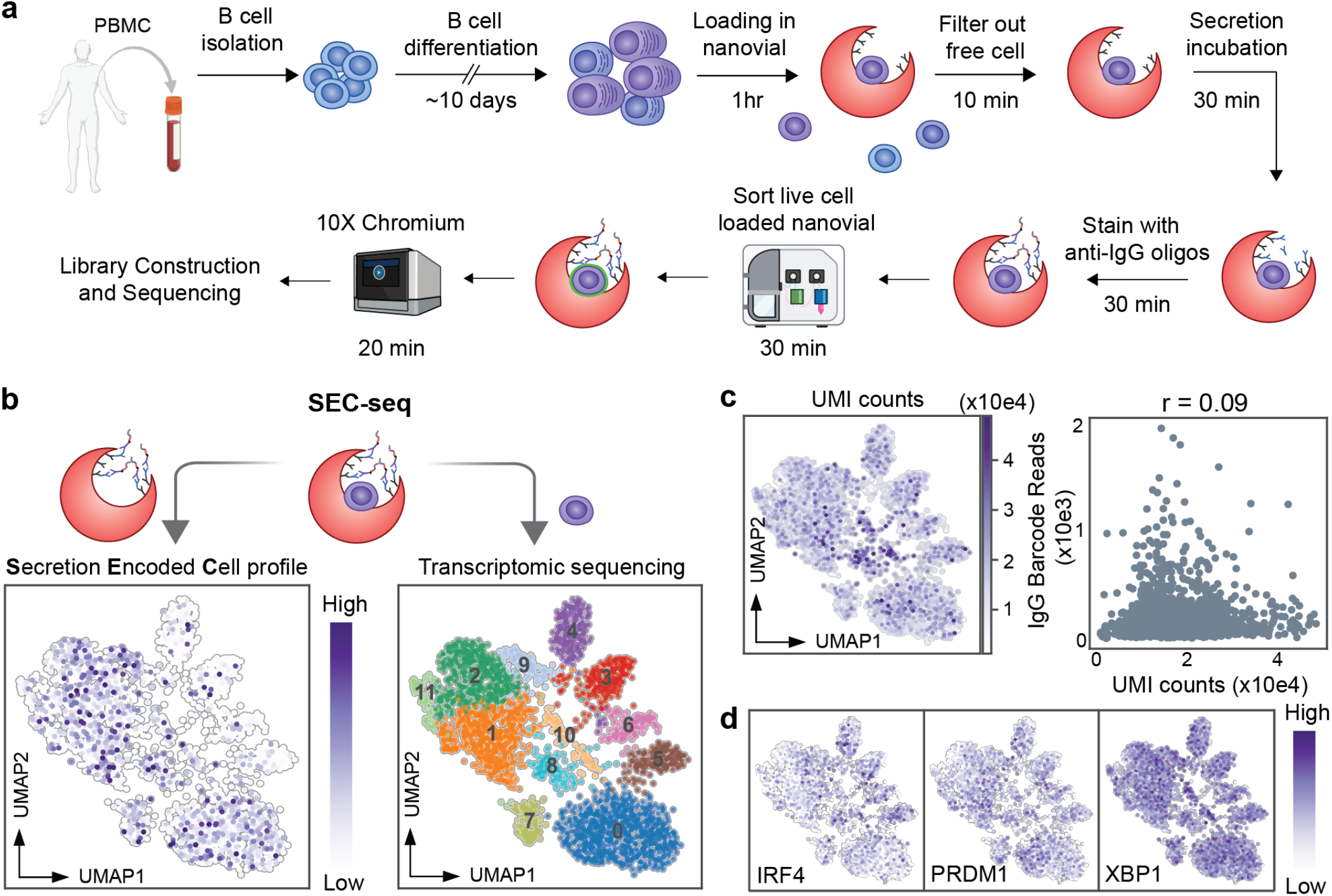
SEC-seq: single-cell transcriptomic sequencing associated with IgG secretion. **(a)** Workflow for SEC-seq to link IgG secretion to transcriptome at the single-cell level. **(b)** Transcriptome-based clustering of single-cell expression profiles of human ASCs (right), and corresponding secreted IgG levels projected on the same UMAP plot (left), n = 3060. **(c)** UMI counts projected on the UMAP (left), and scatter plot of UMI counts and IgG barcode reads (right). r is Pearson correlation coefficient. **(d)** Transcript levels of representative genes projected on the UMAP plot from panel (b) represents transcription factors (TFs) for plasma cells (PCs).

### Using SEC-seq to determine transcriptional signatures associated with IgG secretion

To classify PCs by isotypes, we used a similar strategy as we previously used to gate double negative cells by flow cytometry. The distributions for the gene counts of IgA and IgM were bi-modal (Fig. 5a). We drew “gates” for *IGHM*^*+*^, *IGHA*^*+*^ at the local minimum between the modes in each distribution and further analyzed the DN cells (Fig. 5a). In agreement with the flow cytometry data, we observed a log increase in the median number of unique IgG barcode reads in the DN population relative to the IgM/IgA populations (Fig. 5b). Based on the distribution of IgG barcodes in the *IGHM/A*^+^ cells, we calculated confidence intervals and established a cutoff of 5 logs (∼90% confidence, red line Fig. 5b-c). We then segmented the population by defining IgG secretors as DN cells with unique IgG barcodes exceeding 5 logs plus ∼30% and non-secretors with those with unique IgG barcodes fewer than 5 logs minus ∼30% (dotted lines, Fig. 5c). Approximately 12% of the loaded nanovials were IgG positive in the cell sort (Fig. S8a, right panel), which is comparable to the ∼14% of IgG secretors we quantified in the scRNA sequencing data.

**Figure 5.**
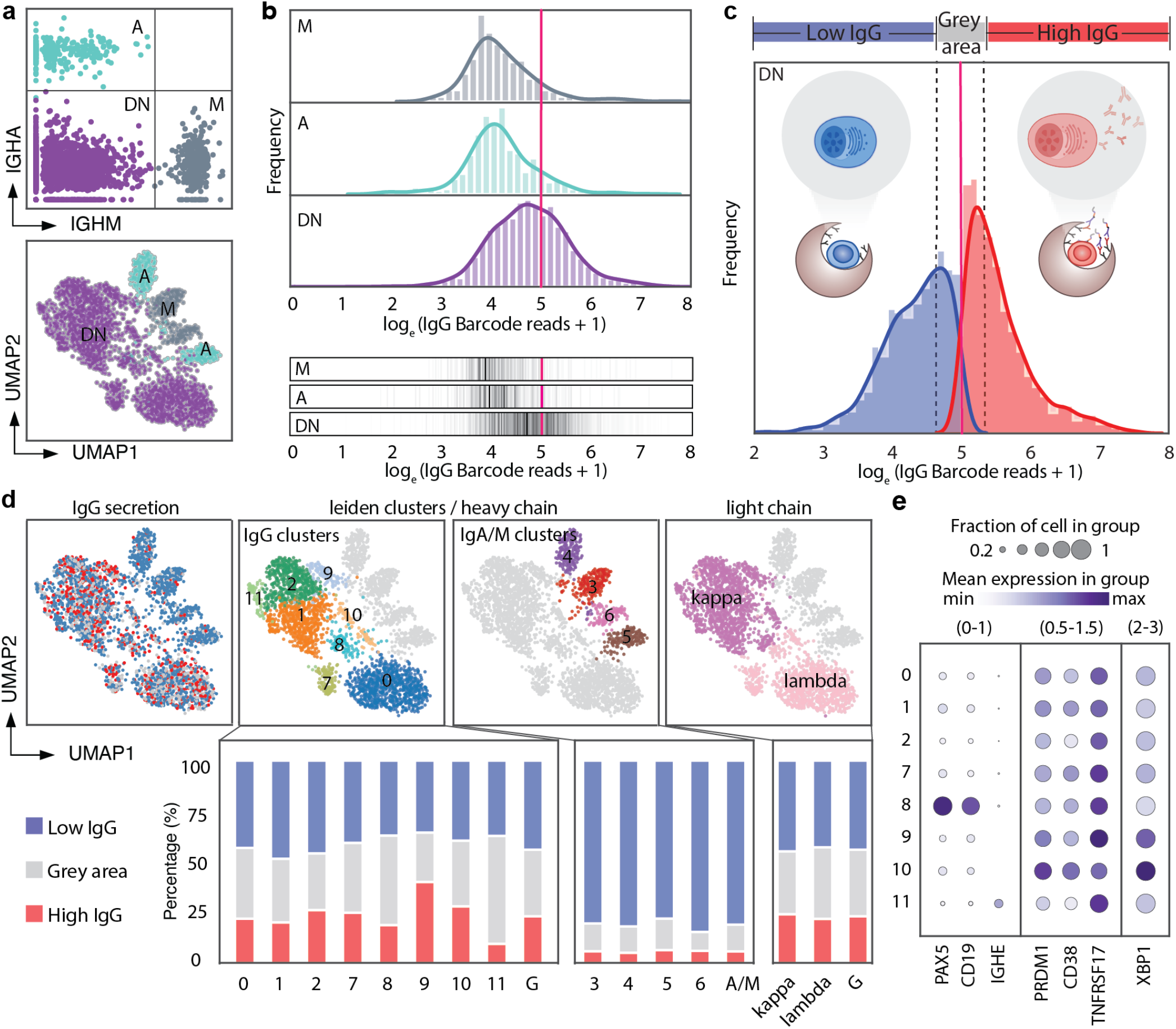
IgG secretors are associated with different gene expression clusters. **(a)** IgM, IgA and DN populations are gated based on expression level of *IGHA* and *IGHM* genes. Identified populations are projected on the gene expression UMAP plot. **(b)** Histogram (upper) and density gradient graph (lower) of unique IgG barcode reads in each subset. Red line is a threshold set at 5 log reads. (**c**) Schematic describing the thresholding used to define high IgG secretors, intermediate (grey area), and non-secretors **(d)** IgG secretors (red) and non-secretors (blue) projected onto UMAP; IgG clusters, IgA/M clusters and light chain clusters of UMAP (upper). Bar graphs display the percentage of in the three thresholded regions using thresholds of below 4.7 (non-secretors), between 4.7 and 5.3 (grey area), and above 5.3 (secretors) for each indicated cluster corresponding to the upper panel. **(e)** Dot plot from numbered clusters identified in (d) with indicated genes associated with a low percentage of IgG secretors (clusters 8,11) and a high percentage IgG secretors (clusters 9, 10). Numbers on top of the dot plot with parentheses indicate the minimum and maximum of mean expression in each column.

Next, we asked how the IgG secretors were distributed in each subcluster. After projection of IgG secretors onto the UMAP (Fig. 5d), we found that the majority of IgG secretors (red dots) overlap with IgG clusters, and a small minority overlap with the IgA/M clusters (see also Fig. S9). Upon looking at the percentage of IgG secretors in each subcluster, we found that IgG clusters have on average ∼ 25% IgG secretors and ∼ 40% non-secretors, whereas the remaining clusters were predominantly non-secretors. The IGHG3 (cluster 9) and IGHG4 (cluster 11) subclusters exhibited higher and lower percentages of IgG-secreting cells, respectively, possibly indicating different levels of IgG secretion among PCs expressing these isotypes. In contrast, we found there was a similar percentage of IgG secretors and non-secretors when cells expressed either light chain kappa or lambda (Fig. 5d). One subcluster within the IgG group expressed the PC progenitor markers *PAX5* and *CD19* but exhibited a similar percentage of IgG-secretors (cluster 8, Fig. 5d-e), demonstrating that a subset of PC progenitors are capable of IgG secretion. Finally, although the PC markers *CD38, PRDM1* (also known as BLIMP1), and *TNFRSF17* (also known as BCMA) do not vary greatly among the subclusters, *XBP1 (*a master regulator of the unfolded protein response) is enriched in clusters 9 and 10,^11^ those that have the largest numbers of IgG-secreting cells (Fig. 5d-e).

### IgG secretion is highly regulated by mitochondrial metabolism and protein transport

Identifying genes associated with IgG secretion is a primary objective of this work. We conducted differential gene expression analysis to compare IgG secretors and non-secretors (see definition above). We then applied gene enrichment analysis (GSEA) to GO: biological process (Fig. 6a). The most highly enriched genes were related to mitochondrial associated gene sets, including ATP synthesis coupled electron transport, oxidative phosphorylation, mitochondrial electron transport NADH to ubiquinone (Fig. 6a, Fig. S10a), etc. The other major enriched genes are translation process and trafficking proteins, e.g. endoplasmic reticulum (ER) to golgi vesicle mediated transport and protein localization to ER (Fig. 6a, Fig. S10a). We also applied GSEA to the HALLMARK dataset,and found MYC-target genes are highly enriched (Fig. 6b, Fig. S10b). Previous studies demonstrated that Myc target genes are highly associated with ribosome function, oxidative phosphorylation, protein export, etc^12,13^.

**Figure 6.**
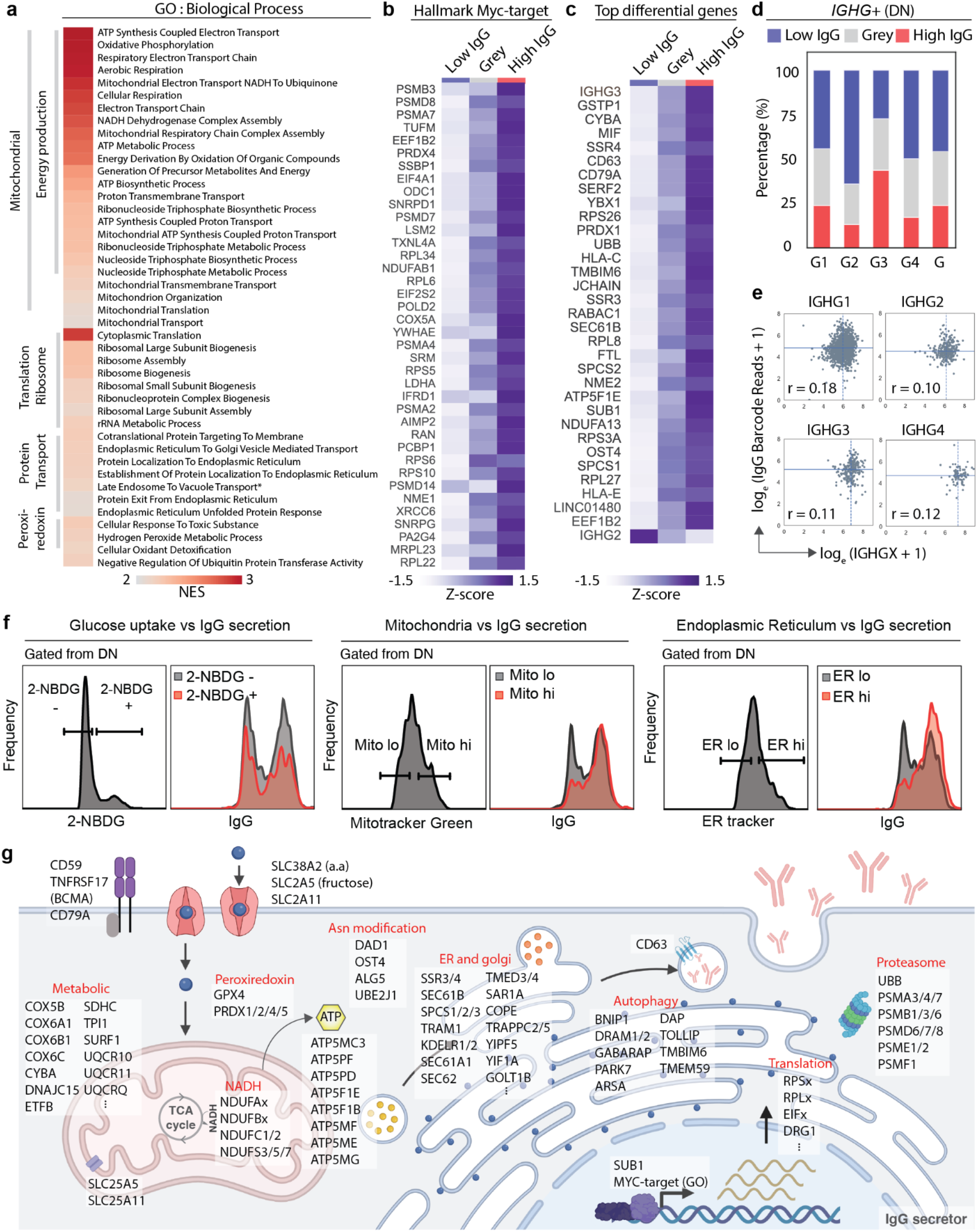
Analysis of SEC-seq data to identify enriched genes associated with IgG secretion. **(a)** Top 40 Gene set enriched list in GO biological process from gene set enrichment analysis (GSEA); heatmap represents the normalized enrichment score (NES). **(b) ∼**Top 40 genes enriched in HALLMARK MYC TARGETS V1 from GSEA. Heatmap represents the z-score of mean of gene expression from each group. **(c)** Top differentially expressed genes and highly expressed (adjusted p-value < 0.1 and mean of gene expression > 5) of IgG secretors. Heatmap represents the z-score of mean of gene expression from each group.**(d)** Bar graphs display the percentage of cells in the three thresholded regions using thresholds of below 4.7 (non-secretors), between 4.7 and 5.3 (grey area), and above 5.3 (secretors) for each sub-class of IgG. **(e)** Scatter plots of IGHG log transcripts and log IgG barcode reads. The horizontal lines represent the median of IgG barcode reads and the vertical lines represent the median of IGHG transcript reads. r is the Pearson correlation coefficient. **(f)** Histograms of IgG secretion levels compared with glucose uptake +/-, high/low mitochondria activity and high/low ER amount. (**g**) Schematic cartoon of top 500 genes upregulated in IgG secretors based on different gene function clustering (DAVID) and GSEA.

We then listed the top highly and differentially expressed genes between secretors and non-secretors (Fig. 6c). Several UPR-relevant genes, e.g. SSR3, SSR4, and SEC61B, are known for being highly expressed in PCs^11,14^. Among the IgG isotypes, *IGHG3* transcripts were increased in IgG secretors, although *IGHG2* transcripts were decreased. As expected PCs expressed only one antibody isotype and the majority of IgG cells expressed IgG1 (75%), followed by IgG2 (12%), IgG3 (9%), and IgG4 (3%). As implied by the cluster analysis we found that *IGHG3-*expressing PCs had a higher percentage of IgG secretors, whereas those expressing *IGHG2* and *IGHG4* had fewer (Fig. 6d). Regardless of the isotype, we found little correlation between steady-state IgG transcripts and IgG secreted protein (Fig. 6e), which implies that steady-state transcript levels do not predict the degree of antibody secretion in PCs.

Contrary to our expectation, we found that transporters of fructose (SLC2A5), but not glucose (e.g. SLC2A1) were expressed more highly in IgG secretors. We evaluated glucose import by incubating PCs with the glucose analogue 2-NBDG. Consistent with the idea that the levels of glucose import do not drive antibody secretion we found that within DN cells there is no association between 2-NBDG import and IgG secretion (Fig. 6f left). In contrast, after staining for mitotracker and ER, we found that PCs with high mitochondrial volumes or cells with high-ER content were predominantly IgG secretors (Fig. 6f, middle panel and right panel). Based on this integration of single-cell secretion and gene expression data, it is likely that the rate-limiting determinants of antibody production/secretion are the cellular programs required for protein secretion, rather than transcript availability. These programs include those regulating protein translation, ER/golgi transport, and protein post translational modification accompanied by mitochondrial respiration, autophagy, and oxidant detoxification (Fig. 6g).

## DISCUSSION

The new methods introduced in this work allowed us to simultaneously interrogate the degree of IgG secretion with surface markers and mRNA at the single-cell level in primary human B cell subpopulations. In single experiments, we were able to analyze more than 3000 cells, directly linking IgG secretion amount with transcriptomes, and more than 8000 cells linking IgG secretion to surface markers. This unprecedented scale of experiments enabled characterization of secretion phenotypes in subpopulations of PCs. PCs that exclusively expressed *IGHG3* and to a lesser extent *IGHG1* were found to have a larger fraction of the subpopulation with IgG secretion signal compared to other subclasses. This could reflect a differential secretion rate or differences in anti-IgG binding that is subclass dependent^15,16^. Importantly, our data reinforce the need to assay cell secretory function directly, since gene expression of the underlying protein is not highly correlated to protein levels^17^ or protein secretion. For example, the level of expression of any of the IgG heavy chain subclasses (IGHG1-4) individually had poor correlation to IgG secretion (Fig. 6e, r=0.10-0.18).

By leveraging standard equipment and workflows, the SEC-seq approach should be widely adoptable by other researchers, which promises to amplify the impact of the technique. Cells loaded on nanovials can be directly input into the microfluidic droplet generator used in the 10X Chromium system, and all other processes follow standard workflows for single-cell transcriptomics. Nanovials do not interfere with lysis, reverse transcription, or downstream cDNA library preparation or sequencing steps. The number of transcripts and gene count for cells on and off of nanovials remained similar (Fig. 3b) and doublet events remained comparable to those observed in cells alone (Fig. 3d). In fact, all of the components used to perform SEC-seq are commercially available including the biotinylated nanovials (Partillion Bioscience) and oligonucleotide barcoded antibodies (Biolegend). Linking secreted protein function with specific nucleic acid sequences of heavy and light chains of IgG (Fig. 3d) or alpha and beta chains of T cell receptors (TCRs) can enable functional discovery workflows for new monoclonal antibodies or engineered TCR-based therapies^6^. A parallel study is also exploring the transcriptome that underlies secretion of high levels of vascular endothelial growth factor by mesenchymal stromal cells (unpublished work), which can elucidate features associated with therapeutic sub-populations of cells. We demonstrate here that the technique is compatible with standard CITE-seq^18^ reagents to create workflows that link surface proteins, secreted proteins, and single-cell transcriptomes^19^. More broadly, nanovials can link surface markers and secreted proteins in viable cells selected using multicolor FACS. Here we demonstrated an 8-color panel, but like surface-marker only panels, the scale of markers should only be limited by the spectral overlap of fluorophores.

CD138 has long been identified as a marker of differentiated human PCs, including long-lived PCs^20,21^ that functionally regulates plasma cell survival in the bone marrow^22,23^. However, there is no reported association between CD138 surface expression with the degree of antibody secretion. Here, we provide definitive evidence that CD138 is a useful marker for *ex vivo* differentiated human PCs that are actively secreting IgG. The marker enriches for the largest fraction of IgG secretors (>75%) compared to other markers like CD38. Similar to what has been observed previously^24,25^, we also found that a subset of CD38low cells secrete IgG, and do so at a slightly decreased degree than other IgG secretors. In addition to the biological implications of these findings, the study also suggests that future work to isolate antibody sequence information from PCs, e.g. for discovery of monoclonal antibodies^26^, should have the most success using CD138 as a marker for magnetic or nanovial-based enrichment.

Antibody-secreting plasma cells exhibit unique biochemical features that enable prolific protein secretion (reviewed in^27,28^). Protein secretion by PCs is regulated by a transcriptional program dependent on PRDM1 (also known as Blimp1) and the unfolded protein response gene, XBP1. PRDM1 is a multifunctional transcriptional activator and repressor involved in the maturation of developing B cells into PCs^29,30^, and in maintenance of immunoglobulin secretion by PCs^31^. As expected, we observed that high IgG secretors exhibited increased mitochondrial and ER volume, as well as transcription of core pathways regulating mitochondrial biogenesis, ER, translation and processing of antibodies. These core secretory pathways are regulated by mTOR, which is activated by PRDM1^32^ in PCs. We also identified several unexpected genes that are increased in IgG secretors and may be critical for their function including CD79A, the complement regulator CD59 and macrophage inhibitory factor MIF, which could regulate cells in the PC bone marrow microenvironment.

XBP1 is most highly expressed in the clusters with highest IgG secretion, which suggests that increased levels of the unfolded protein response pathway may facilitate increased immunoglobulin secretion^33^. In contrast, PC-specific knockout of XBP1 eliminates immunoglobulin secretion^31^ by PCs without eliciting cell death. Collectively, these results suggest that tuning XBP1 levels could transiently increase or decrease the amount of antibody secreted by PCs. To be activated, XBP1 transcripts are regulated at the level of splicing by the upstream unfolded protein response gene IRE1^34^. Recently, pharmacological compounds have been developed that increase^35^ or decrease^36^ IRE1 activity, and thus regulate the degree of spliced, active XBP1. In future studies, we envision using nanovials and/or SEC-seq to identify nontoxic pharmacological and/or genetic strategies that can increase and/or decrease immunoglobulin secretion. We envision that such molecules could be used to transiently increase the antibody responses to pathogens and/or decrease antibody responses in antibody-driven disease like lupus or arthritis.

We show that nanovial technology and SEC-seq can be used to link the levels of antibody secretion to cell surface markers, transcriptional signatures, and vital dyes. This technology enables us, for the first time, to study the molecular determinants of protein secretion by PCs. Because high and low secreting cells can be captured by cell sorting, we predict that this technique will be used to study PC secretion at a single cell level and co-evaluate a myriad of parameters including, but not limited to epigenetics, metabolomics, signaling, loss of function and mutational scans. Beyond the analysis of secreted immunoglobulins, the technique can unlock the ability to study >3000 proteins that are part of the human secretome,^37^ ultimately identifying shared, or unique, molecular underpinnings of secretion pathways that are critical for cell communication and function from the single-cell to the organismal level.

## MATERIALS AND METHODS

### Cell culture methods

All cells were cultured in incubators at 37°C and 5% CO2 in static conditions unless otherwise noted.

#### Human primary B cells

We isolated B cells from healthy donors’ peripheral blood mononuclear cells (Fred Hutchinson Cancer Research Center) using the EasySep Human B cell isolation kit (Stem Cell Technologies). Isolated B cells were cultured in Iscove’s modified Dulbecco’s medium (Gibco) supplemented with 2-mercaptoethanol (55 µM) and 10% FBS. Cells were cultured for seven days (activation) in medium containing 100 ng/mL megaCD40L (Enzo Life Science), 1 µg/ml CpG ODN2006 (invitrogen), 40 ng/mL IL-21 (Peprotech), and then for three to five days (plasmablast/plasma cell differentiation) in medium containing 50 ng/mL IL-6 (Peprotech),10 ng/mL IL-15 and 15 ng/mL interferon-*α*2B (Sigma-Aldrich).

#### Hybridoma cells

HyHel-5 cells were maintained in IMDM media (Invitrogen 12440053) supplemented with 10% FBS (Invitrogen 16000044) and 1% penicillin/streptomycin (Invitrogen). Cells were passaged down to a final concentration of 2×10^5^ cells/mL every three days.

#### Raji cell

Cells were sourced from ATCC (CCL-86™) and maintained in RPMI-1640 ATCC modification media (Invitrogen A1049101) supplemented with 10% FBS (Invitrogen) and 1% penicillin/streptomycin (Invitrogen). Cells were passaged down to a final concentration of 2×10^5^ cells/mL every three days.

### Methods of coating nanovials

Nanovials were coated with streptavidin by mixing equal volumes of streptavidin (300 *µ*g/mL) and nanovials (n=340,000) for 15 mins at room temperature followed by three washes which consisted of resuspension in clean washing buffer (supplementary Table S1) and centrifugation at 400 x g. The coating antibody mix was prepared by making a 10X dilution of biotinylated anti-IgG (0.5 mg/mL) and biotinylated anti-cell surface protein (anti-CD27 or anti-CD45) (0.5 mg/mL) in washing buffer (final antibody concentration 0.05 mg/mL per antibody). Equal volumes of the coating antibody mix, and the streptavidin coated nanovials were combined and placed at room temperature for 30 minutes or 4°C overnight. Antibody coated nanovials were washed twice in washing buffer. After the final wash, the supernatant was removed and the nanovials were resuspended in cell media. The final prepared nanovials were placed on ice until cell loading.

### Methods of cell loading in nanovials

Differentiated human B cells were first processed by ficoll density gradient centrifugation to remove dead cells and debris. For secretion experiments, 5 µL of blocking antibody (anti-IgG, Southern Biotech) was added to the 50 µL cell solution to yield a final concentration of 25 µg/mL. Then 55 µL of the cell solution (244,000 cells) was mixed with 20 µL of concentrated nanovials (n= 340,000) at a ratio of 1:1.4 (cell: nanovial) on ice by pipetting for 30 seconds with a circular motion. Cells were loaded into nanovials by carefully pipetting throughout the pellet of nanovials. We then added 1 mL of biotin-free medium and incubated the mixed cells and nanovials on ice for 1 hour without any perturbation.

### Methods of incubation of cells with nanovials

After the 1 hour loading process, samples were placed on top of the strainer and washed by 1 mL wash buffer two times. A 15 mL falcon tube was precoated with washing buffer, and used for cell collection by inverting the strainer and placing on top of the tube. 2 mL of washing buffer was used to wash the nanovials off the strainer and into the falcon tube. We then centrifuged the 15 mL falcon tube containing the filtered nanovials at 300 x g at 4°C for 5 min, and resuspended nanovials into B cell medium pre-warmed to 37°C. The falcon tubes containing nanovials in media were then incubated at 37°C on a rotator for gentle agitation (10 rpm) for 30 minutes to accumulate secretions on nanovials.

### Methods of cell staining of surface markers and secreted IgG

FACS tubes were precoated with staining buffer (supplementary Table S1). After incubation to capture secretions, nanovial samples in falcon tubes were then centrifuged at 400 x g for 5 mins and resuspended into FACS tubes with a cocktail of antibodies (supplementary Table S2) to stain cell surface markers and secreted IgG in staining buffer. Samples were stained on ice for 20 minutes, then washed twice, before analysis by flow cytometry.

### Methods of flow cytometric analysis

Flow cytometric analysis was performed on an LSR II flow cytometer (BD Biosciences) and events were analyzed using FlowJo software (BD). Flow cytometry gating for fluorescent proteins and viability, and immunophenotyping can be found in the supplement.

### Methods of imaging flow cytometric analysis

An ISX 493 MKII equipped with a 405, 488, 560, 592, 642 and 785 nm lasers were used to interrogate B cell secretion capture on nanovials. 40000 objects were collected under 20x magnification with laser power and channel assignments as listed in Supplemental Tables S3 and S4.

Quantification of IgG signal on nanovials from cell secretions was performed through the gating strategy illustrated in Supplemental Figure S4. First, objects in focus were identified by the GradientRMS function in the brightfield channel. Single nanovial objects were separated from debris and multicellular aggregates based on an Aspect Ratio vs Area. Visual inspection of objects located in the gates drawn in S4B confirms correct placement of gate settings. CD38/CD138 double positive cells were further identified via the intensity feature on the CD38 and CD138 channel and separated into high and low signal populations. The amount of IgG secretion for these four populations was quantified by constructing a “nanovial mask” by subtracting the area localized to cells (CD38 or CD138) from the mask identified in the brightfield channel.

### Methods for visualizing nanovials through 10X Genomics Chip G

Fluorinated oil with surfactant, gel bead solution and nanovial solutions were added into the reservoirs of the 10X Genomics Chip G. 3 mL syringes (Becton Dickinson) were connected to the bead and sample inlet reservoirs via PEEK tubing (IDEX) and a coupler molded out of PDMS. Syringe pumps (PhD 2000, Harvard Apparatus) were used to inject air into the reservoirs and pressurize the bead and sample inlets and drive flow. Droplet formation videos were recorded using an inverted microscope (Nikon TE300) equipped with a high-speed camera (ZWO ASI144MM).

### Mixed species SEC-seq validation experiment

Barcoded nanovial preparation (Raji sample). Nanovials (4 million/mL) were modified by mixing a solution of nanovials in washing buffer with a cocktail of 40 µg/mL streptavidin and 20 µg/mL TotalSeqC conjugated streptavidin (Biolegend) solution at equal volumes. The sample was then placed on a rotator (speed) at room temperature for 30 minutes. Nanovials were washed three times by centrifuging the sample at 200 xg for 5 minutes, removing the supernatant, and resuspending in washing buffer. Following washing, the streptavidin modified nanovial solution was then mixed with a solution containing 8 μg/mL anti-human CD45 (Biolegend) at equal volume. The nanovials were then placed on a rotator at 4°C overnight. The nanovials were then washed three times as above and resuspended in cell media before proceeding with cell experiments.

Non-barcoded nanovial preparation (Hybridoma sample). Nanovials (4 million/mL) were modified by mixing with a solution of 60 µg/mL streptavidin at equal volume following standard procedures as described above. The streptavidin modified nanovial solution was then mixed with a solution containing an antibody cocktail comprising 8 µg/mL anti-mouse CD45 (R&D Systems) biotinylated according to manufacturer instructions. (Thermo) and 20 µg/mL Goat anti-Mouse IgG FC (Jackson Immuno Research) at equal volume.

Raji and Hybridoma cells were loaded separately into nanovials on ice and incubated for 1 hour to allow binding. Background cells were removed using a 20 μm cell strainer and cells were then incubated in a CO2 incubator at 37°C for 30 min to accumulate secreted IgG. The hybridoma sample was stained with a 12 μg/mL solution of TotalSeqC conjugated anti IgG1 antibody (Biolegend) and incubated for 45 minutes on Ice. Samples were then washed three times with ultra-pure PBS containing 0.04% BSA (Invitrogen). The Raji and Hybridoma samples were mixed at a 1:1 ratio prior to proceeding with single-cell sequencing library preparation. Libraries were prepared at the UCLA sequencing core using the 10X Genomics Chromium Next GEM Single Cell 5’ Kit v2 + Feature barcode libraries. Approximately 8000 nanovials were added in the sample lane and emulsified with barcoded beads with the Chromium instrument. Libraries were QC’d using the tapestation (Agilient) and sequenced using NovaSeq S2 (100 Cycles). FASTQ files were processed by cellranger and the Barnyard reference genome (refdata-gex-GRCh38-and-mm10-2020-A) was used for alignment. An additional custom reference genome was appeneded for alignment to identify transcripts for the known heavy and light chain sequences of the hybridoma cell line (HyHel-5). Data visualization and analyses was performed using Loupe Browser (v6.0) and Seurat.

### SEC-seq human ASCs sample preparation and sequencing

Following *ex vivo* differentiation, cells were treated with Ficoll to remove any debris. Viable cells were immediately loaded onto nanovial and incubated to accumulate secretions, as described above. After accumulating secretions on the nanovials, samples are stained with PE anti-IgG (BD), and a secondary antibody (anti-PE) that was labeled with barcoded oligonucleotides (Biolegend). Next, we sorted viable cells loaded on nanovials with the WOLF cell sorter (Nanocellect) by threshold on the negative SYTOX population (Figure S8). Sorted cell-loaded nanovials were introduced into the 10X Genomics Chip G (10X Genomics), at 2500-10000 cell-loaded nanovials per lane. Next, we prepared libraries using 10X Genomics Chromium Next GEM Single Cell v3.1 kit following the 10X user guide (CG000317). Libraries from the oligonucleotide barcoded antibodies bound to nanovials and transcripts were evaluated by tapestation (Agilent) before sequencing. Finally, libraries were pooled at a ratio of 80% cellular RNA to 20% oligonucleotide barcoded antibodies and sequenced with NextSeq 1000/2000 kit (Illumina) using the following read length: 28 bp Read1, 10 bp i7 Index, 10 bp i5 Index, and 90 bp Read2.

### Single Cell RNA-seq analysis

FASTQ files were processed by cellranger based on the human reference genome GRCh38. The h5 file was then further analyzed using a custom python script. Data analysis including normalized, dimensional reduction, hierarchical clustering, leiden clustering and differential gene analysis were performed with scanpy^38^.

### ELISpot (enzyme-linked immunospot)

PBS pre-wetted 96 well plates (Millipore) were coated overnight at 4 °C with goat anti-human IgG (H + L) capture antibody (Jackson Immunoresearch). 100 µL of IMDM was used to block each well for 2 hours at 37°C. A designated number of cells were washed with PBS and resuspended in 2X cytokine cocktail in IMDM. We added 100 µLof cells to the elispot plate directly and incubated at 37°C in an incubator overnight. Cells were then washed away, and plates were washed six times. Secreted IgG was detected by binding to HRP-conjugated goat anti-human IgG secondary antibody (Southern Biotech). Spots were developed with AEC Substrate Kit, Peroxidase (HRP) reagents (Vector Laboratories). The spots were quantified using the Cellular Technology Limited; CTL ImmunoSpot software.

## ACKNOWLEDGEMENTS

We would like to thank other members of the James lab, Partillion Bioscience and Di Carlo lab for helpful comments and discussion in preparation of this manuscript. We thank 10X Genomics for providing reagents for these studies. We thank Nanocellect for providing access to a WOLF Sorter to perform sorting, and Donna Munoz for her assistance in these efforts. We would like to thank the Brotman Baty Institute and the UCLA Technology Center for Genomics & Bioinformatics for performing sequencing services. Last, we would like to thank Huiyun Sun in Fred Hutch for offering help for library QC.

## AUTHOR CONTRIBUTIONS

RYC, JD, DD and RGJ designed the study and wrote the manuscript. ARO and RYC cultured human B cells. RYC performed the IgG secretion experiment with flow cytometry and analyzed the data. BEH ran samples on the ImageStream and assisted in downstream analysis of the collected data. JD, DD, and WY designed the initial SEC-seq validation experiments which were performed by LB and WY and analyzed by JL and JD. RYC, JD, DD and RGJ designed the SEC-seq human plasma cell experiment. RYC and ARO performed the SEC-seq experiment. RYC analyzed the SEC-seq data.

