## Supplemental materials for "SEC-seq: Association of molecular signatures with antibody secretion in thousands of single human plasma cells"

**Supplemental Tables**

Table S1. Buffer and reagents
Table S2. Antibodies
Table S3. ImageStreamX® MKII fluorochrome layout
Table S4. ImageStreamX® MKII Collection Information

### Table S1. Buffer and reagents

| Buffer/Medium | Item | Vendor | Catalog | Concentration |
|---|---|---|---|---|
| Washing buffer | DPBS | Fisher Scientific | 14190250 | |
| | BSA | VWR life science | 0332-100G | 0.50% |
| | Pluronic F-127 | Sigma | P2443-250G | 0.05% |
| Staining buffer | DPBS | Fisher Scientific | 14190250 | |
| | FBS | Sigma | F0926-500ML | 2% |
| | Pluronic F-127 | Sigma | P2443-250G | 0.05% |
| Sorting Buffer | PBS | Cytiva | SH30256.01 | |
| | Blocker BAS (10%) in PBS | Thermo Scientific | 37525 | 1% |
| | Pluronic F-127 | Sigma | P2443-250G | 0.05% |
| B cell loading medium | DMEM+GlutaMAX | Gibco | 10569-010 | |
| B cell secretion medium | Lonza BioWhittaker Iscove's Modified Dulbecco's Medium (IMDM) without L-Glutamine | Fisher | BW12726F | |
| | FBS | Sigma | F0926-500ML | 10% |
| | GlutaMAX 100X | Gibco | 35050-061 | 1X |
| | 2-Mercaptoethanol | Fisher | O3446I-100 | 55um |

### Table S2. Antibodies

| Experiment | Item | Vendor | Catalog | Clone |
|---|---|---|---|---|
| human B cell capture antibody | Biotin anti-human CD45 Antibody | Biolegend | 304004 | HI30 |
| human B cell capture antibody | Biotin anti-human CD38 Antibody | Biolegend | 303518 | HIT2 |
| human B cell capture antibody | Biotin anti-human CD27 Antibody | Biolegend | 356426 | M-T271 |
| human B cell capture antibody | Biotin anti-human CD31 Antibody | Biolegend | 536604 | O92E4 |
| human IgG capture antibody/blocking antibody | Goat Anti-Human IgG Fc, Multi-Species SP ads-BIOT | Southern Biotech | 2014-08 | Polyclonal |
| Flow Cytometry | PE-Cy7 anti-human CD19 | Biolegend | 302216 | HIB19 |
| Flow Cytometry/Amnis | PerCP-Cy5.5 anti-human CD38 | BD | BDB551400 | HIT2 |
| Flow Cytometry/Amnis | PE anti-human CD138 | Biolegend | 356504 | MI15 |
| Flow Cytometry/Amnis | PacBlue anti-human IgM | Biolegend | 314514 | MHM-88 |
| Flow Cytometry | APC anti-human IgM | BD | 551062 | G20-127 |
| Flow Cytometry | APC-Vio® 770 IgA Antibody, anti-human | Miltenyi | 130-113-473 | IS11-8E10 |
| Flow Cytometry | AF700 anti-human IgG | BD | 561296 | G18-145 |
| Amnis | APC anti-human IgG | BD | 562025 | G18-145 |
| SEC-seq cross-sp | TotalSeq™-C0971 Streptavidin | Biolegend | 405271 | |
| SEC-seq cross-sp | anti-mouse CD45 | R&D Systems | AF114 | Polyclonal |
| SEC-seq cross-sp | Goat anti-Mouse IgG FC | Jackson Immuno Research | 115-065-071 | Polyclonal |
| SEC-seq cross-sp | TotalSeq™-C1167 anti-mouse IgG1 Antibody | Biolegend | 406636 | RMG1-1 |
| SEC-seq | PE Mouse Anti-Human IgG | BD | 555787 | G18-145 |
| SEC-seq | TotalSeq™-B0911 anti-phycoerythrin (PE) Antibody | Biolegend | 408113 | PE001 |

**Table S3. ImageStreamX® MKII fluorochrome layout**

| Ch | Band | Fluorochrome | Target | Notes |
|---|---|---|---|---|
| **1** | **430-470** | Brightfield | Morphology | **FIXED** |
| **2** | **470-560** | | | |
| **3** | **560-595** | | | |
| **4** | **595-660** | PE | CD138 | Plasma cell marker |
| **5** | **660-745** | PerCP Cy5.5 | CD38 | Plasmablast/plasma cell marker |
| **6** | **745-785** | SSC | Granularity | |
| **7** | **430-505** | Pac Blue | IgM | Surface bound, not captured on nanovial |
| **8** | **505-575** | | | |
| **9** | **575-595** | Brightfield | Morphology | **FIXED** |
| **10** | **595-660** | | | |
| **11** | **660-720** | APC | IgG | Antibody released to nanovial by plasma B cells |
| **12** | **720-785** | | | |

**Table S4. ImageStreamX® MKII Collection Information**

| | | | | |
|---|---|---|---|---|
| **Instrument**: ISX 493 | **405 power**: 120mw | **488 power**: 400mW | **560 power**: Off | |
| **592 power:** Off | **642 power:** 150mW | **785 power: 2** mW | **Mag:** 20x | |
| **INSPIRE**: Current | **Bin**: None | **Core diameter**: 10 | **Velocity**: 66 | **%Bd**: 7 |
| **CameraSync**: 38.9 | **Focus**: 0.89 | **Core Tracking**: 11 | **Focus Tracking**: 0 | |

**Supplemental Figures**

Figs. S1 to S10

Figure S1. Nanovial secretion assay gating strategy on flow cytometers and characterization of cell loading using antibodies against different cell surface markers.
Figure S2. ELISPOT spot analysis and IgG secretion heterogeneity.
Figure S3. Nanovial secretion with or without anti-IgG blocking.
Figure S4. Nanovial secretion assay gating strategy on the ImageStreamX MKII Gating Strategy.
Figure S5. Additional images of cell loaded nanovials taken with the ImageStreamX® MKII.
Figure S6. High-speed microscopy images of the gel beads and nanovials loading into droplets using the 10X-Chromium chip G.
Figure S7. Statistics on multiplet events from the mixed species experiments shown in Figure 3.
Figure S8. Sec-seq live cell loaded nanovial sorting.
Figure S9. UMAP of immunoglobulin.
Figure S10. GSEA plot and gene list in core enrichment.

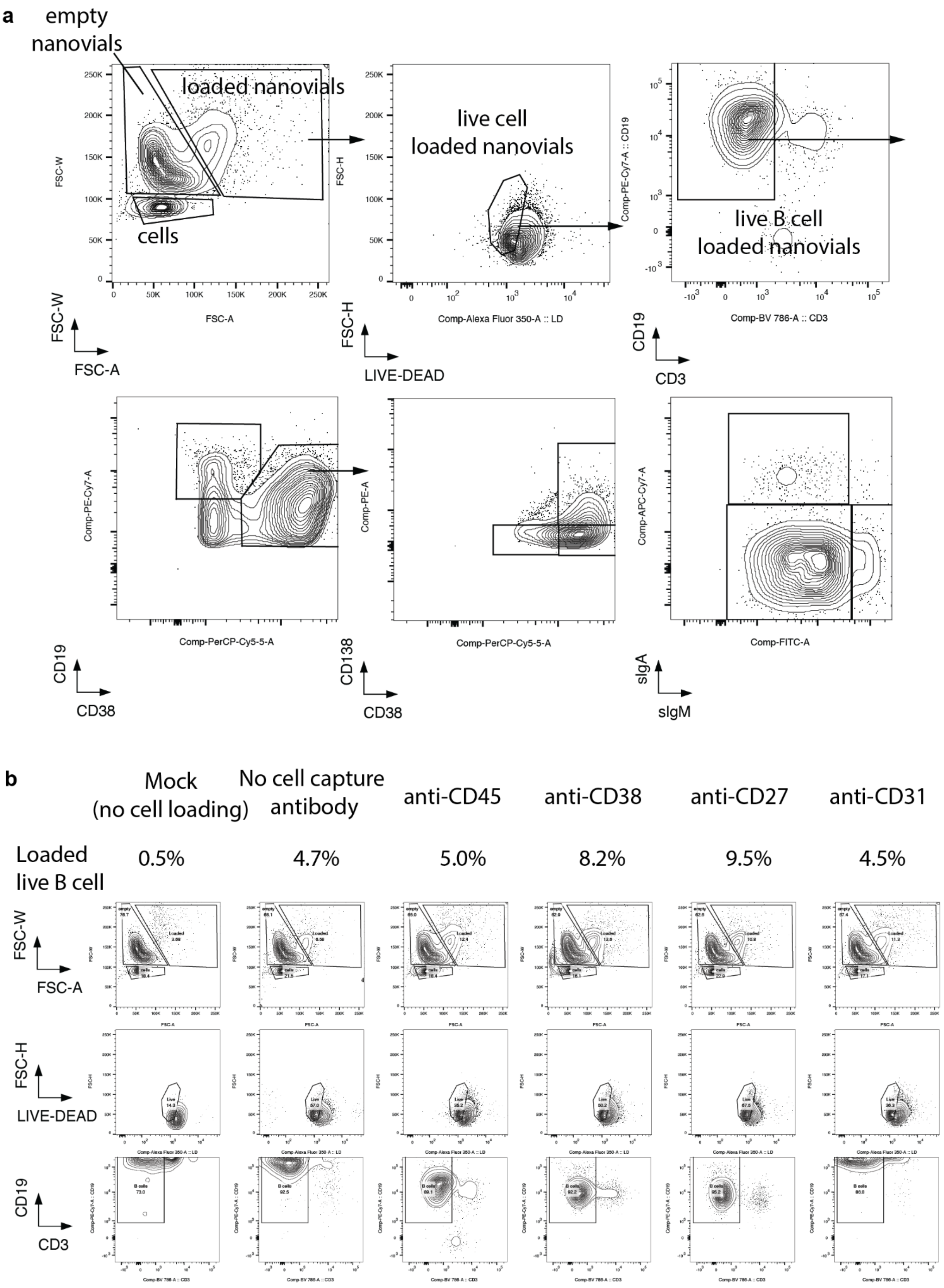
a
empty nanovials
loaded nanovials
cells
FSC-W
FSC-A
live cell loaded nanovials
FSC-H
LIVE-DEAD
Comp-Alexa Fluor 350-A :: LD
live B cell loaded nanovials
Comp-PE-Cy7-A :: CD19
Comp-BV 786-A :: CD3
CD19
CD3
Comp-PE-Cy7-A
Comp-PerCP-Cy5-5-A
CD19
CD38
Comp-PE-A
Comp-PerCP-Cy5-5-A
CD138
CD38
Comp-APC-Cy7-A
Comp-FITC-A
sIgA
sIgM
b
Mock (no cell loading)
No cell capture antibody
anti-CD45
anti-CD38
anti-CD27
anti-CD31
Loaded live B cell
0.5%
4.7%
5.0%
8.2%
9.5%
4.5%
FSC-W
FSC-A
FSC-H
LIVE-DEAD
CD19
CD3

**Figure S1. Nanovial secretion assay flow cytometry gating strategy and characterization of cell loading using antibodies against different cell surface markers.**
**(a)** Gating strategy for live B cell loaded nanovials used for surface marker and IgG secretion analysis. **(b)** Flow plots of cell loading with different cell capture antibodies linked to the nanovials. Loaded live B cell percentage is calculated by dividing the loaded live B cell counts by the total number of nanovial counts (loaded nanovial counts + empty nanovial counts).

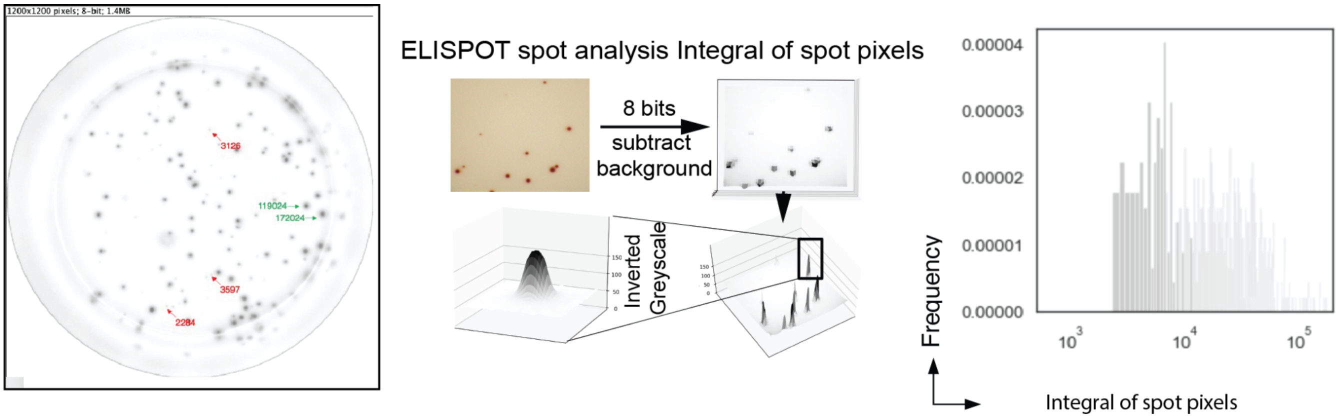


**Figure S2. ELISPOT spot analysis and IgG secretion heterogeneity.**
An ELISPOT image (left) showing IgG secretion for plasma cells following our differentiation protocol. Red arrows point to events with low IgG secretion (small faint spots) and green arrows point to events with higher IgG secretion (larger, more intense spots). The spot intensity is integrated over the entire spot area using ImageJ and the output histogram (right) represents the heterogeneous IgG secretion ability of the population.

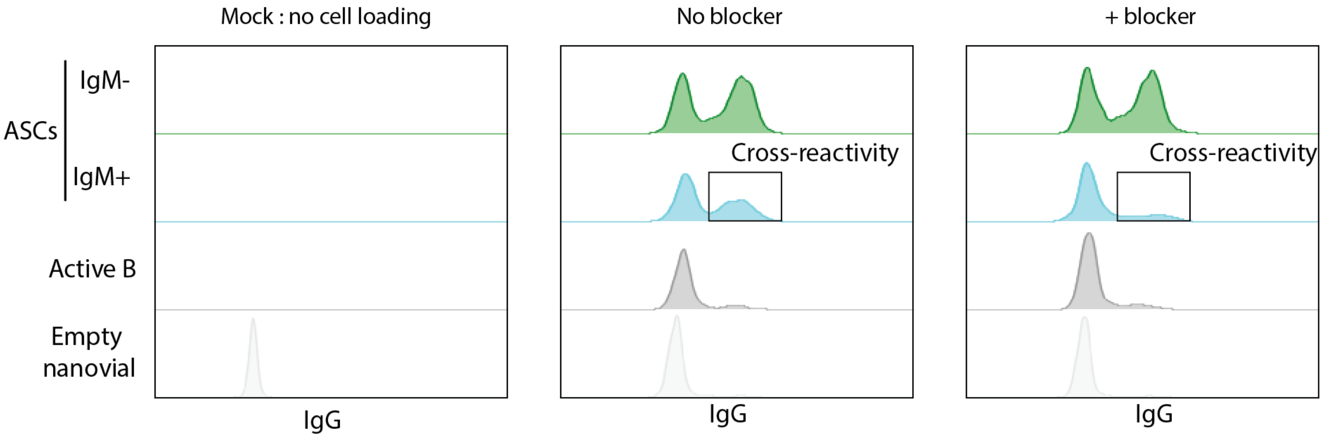


**Figure S3. Nanovial secretion with or without anti-IgG blocking.**
We adjusted the loading strategy to reduce cross-reactive IgG signal to nanovials containing IgM cells. In order to reduce cross-reactivity cells are incubated with anti-IgG as a sink for secreted IgG during the cell loading step and then the anti-IgG solution is washed prior to the cell secretion incubation step.

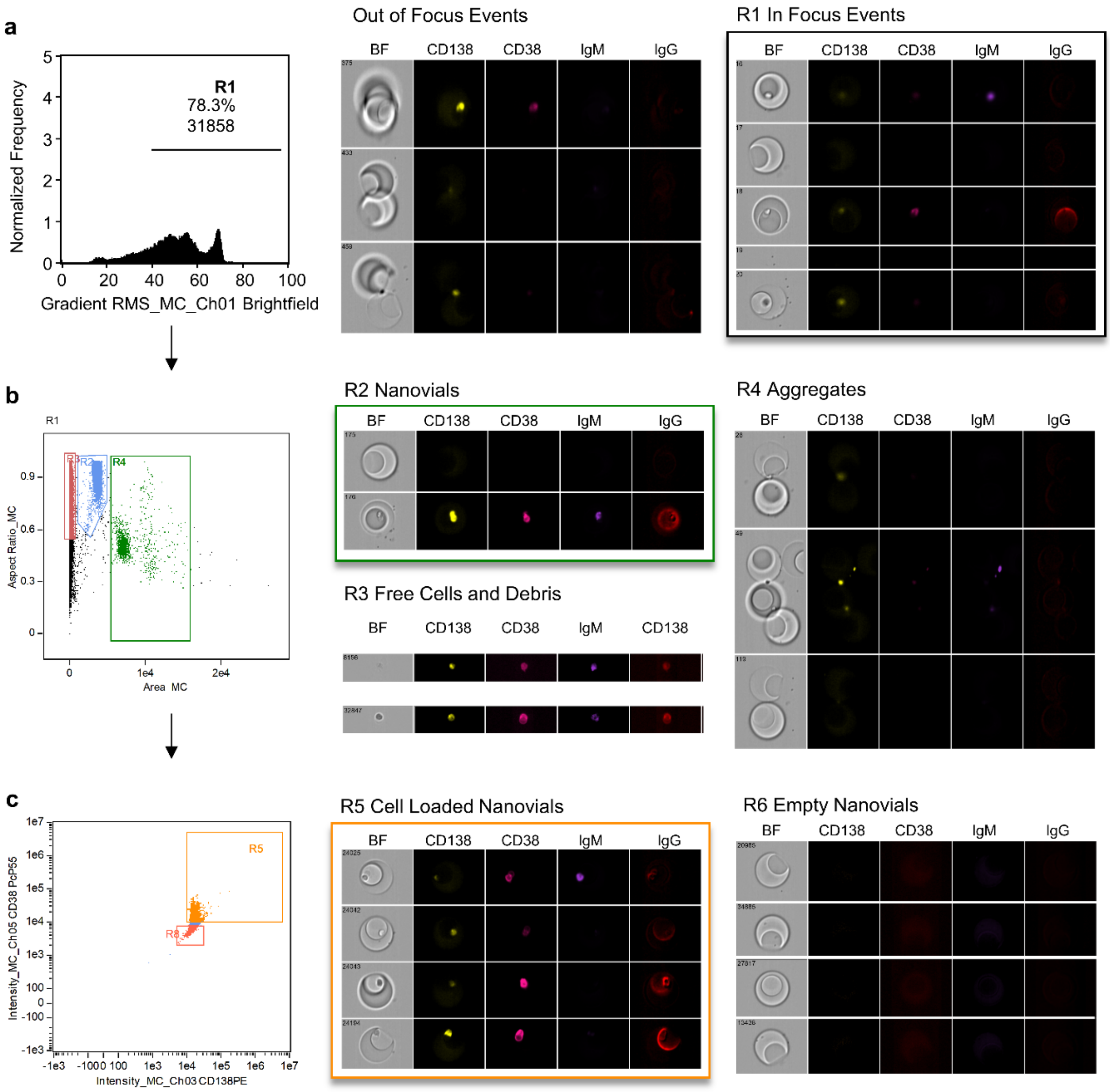


**Figure S4. Nanovial secretion assay gating strategy on the ImageStreamX® MKII. (a)** In focus events are identified by gating on the root mean square gradient of the brightfield channel (Gate R1). **(b)** Events associated with small debris, free cells, nanovials, and nanovial aggregates can be assigned based on the aspect ratio and area of the event. Gate R2 contains single nanovials. **(c)** Cell-loaded nanovials are then gated based on a threshold level of CD38 and CD138 signals (Gate R5).

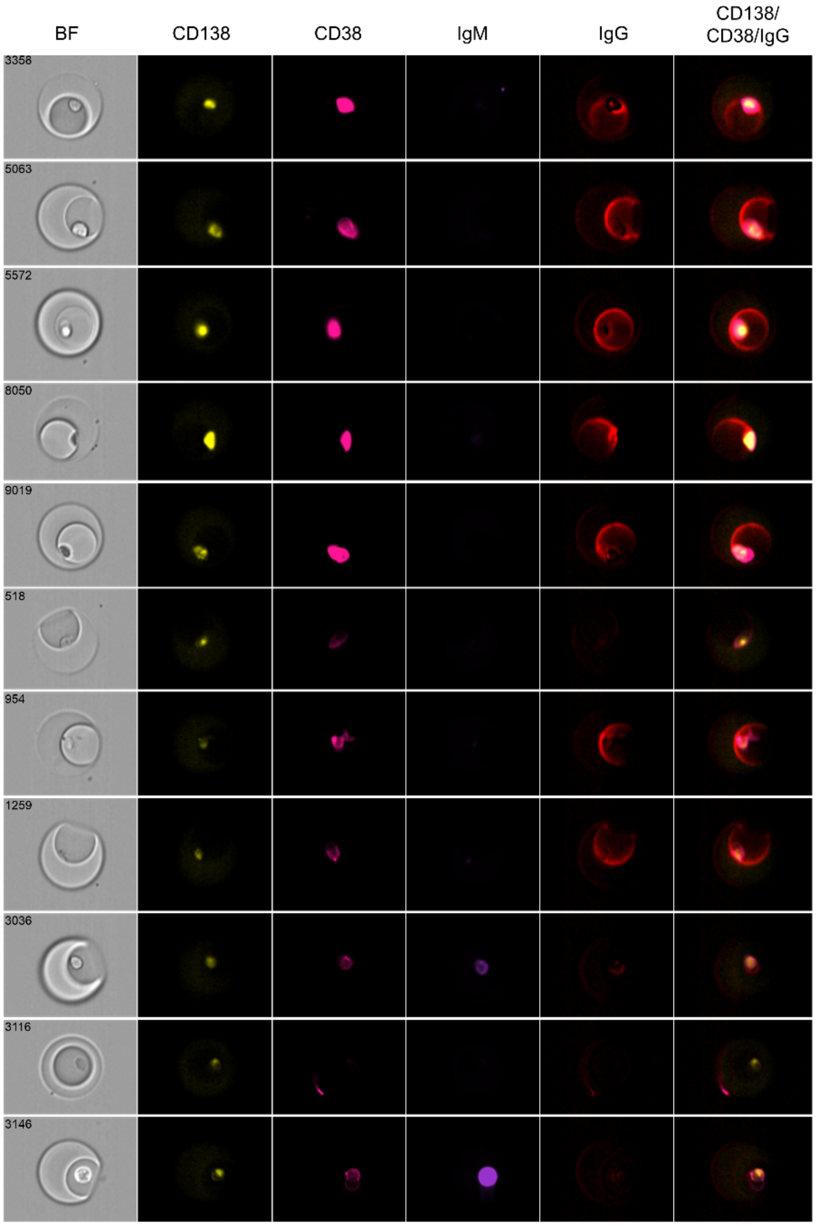


**Figure S5. Additional images of cell loaded nanovials taken with the ImageStreamX® MKII.** Compared to the punctate cell surface signal for surface markers, the secreted IgG signal is clearly seen across the hemispherical cavity of the nanovial.

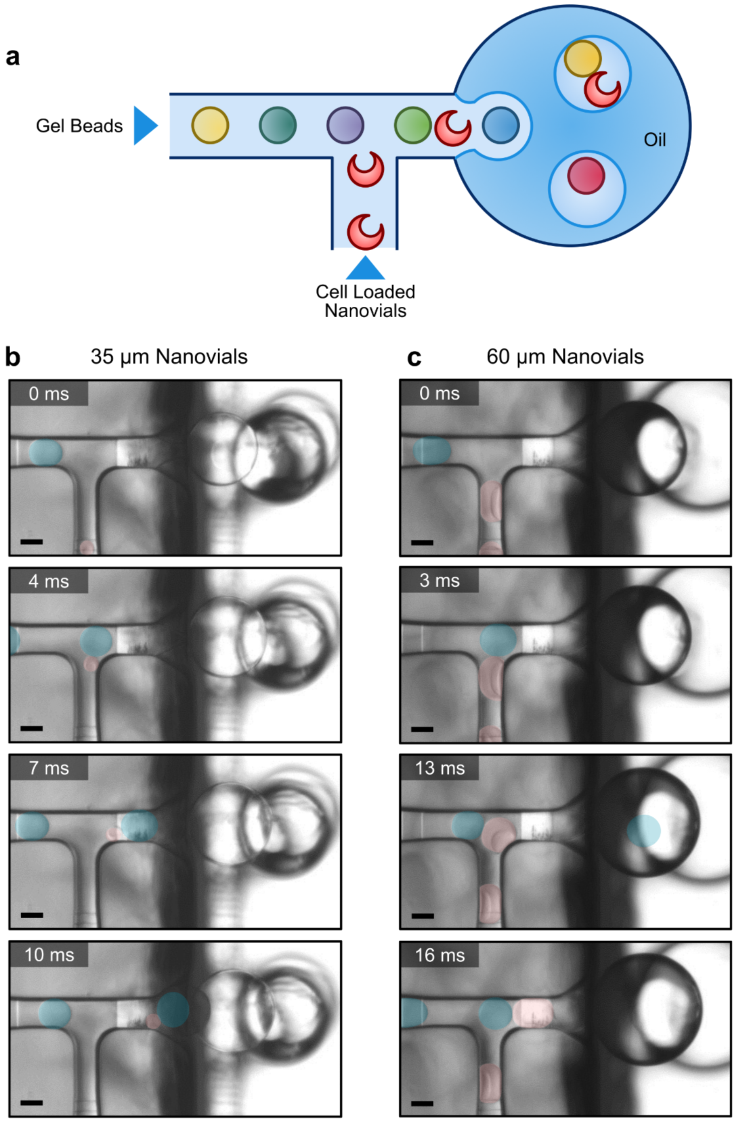


**Figure S6. High-speed microscopy images of the 10X Genomics gel beads and nanovials loading into droplets formed on the 10X Next GEM chip G. (a)** Gel beads and nanovials are loaded through the gel bead inlet and cell sample inlets, respectively, as shown in the illustration. **(b)** 35 µm outer diameter nanovials pass through the cell sample inlet without obstruction and load into forming droplets. **(c)** 60 µm nanovials become constrained in the sample inlet and eventually pass through after deforming. Gel Beads and nanovials are false-colored blue and pink, respectively, to aid in visualization. Scale bar: 50 µm.

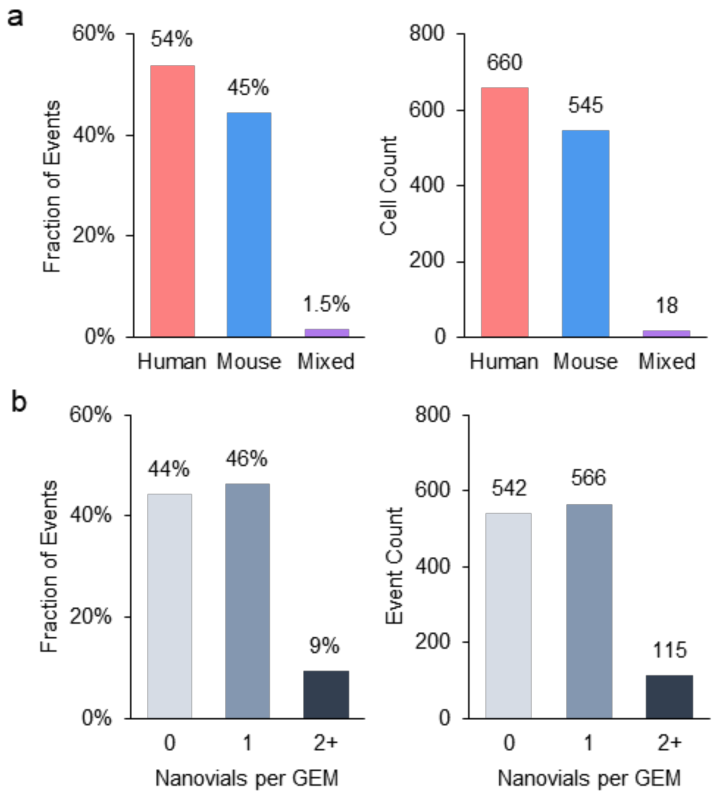


**Figure S7. Statistics on multiplet events from the mixed species experiments shown in Figure 3.** Nanovials were loaded into the 10X Chromium chip at the recommended concentration for cells to achieve 10,000 nanovial events. Approximately 10% of nanovials were loaded with cells and a roughly equal population of cells were off the nanovials. **(a)** The fraction of events based on species classification are plotted. The cell multiplet rate is close to the expected value reported in the 10X literature based on cell count (0.8% expected, 1.5% actual). **(b)** Fraction of events associated with 0, 1, or 2 or more nanovials. Using feature barcode reads on nanovials we discriminated the number of nanovials per cell event. Given there are ~10 times the number of nanovials as cells for this experiment the number of events with multiple nanovials are within the expected range (7.6% expected, 9% actual).

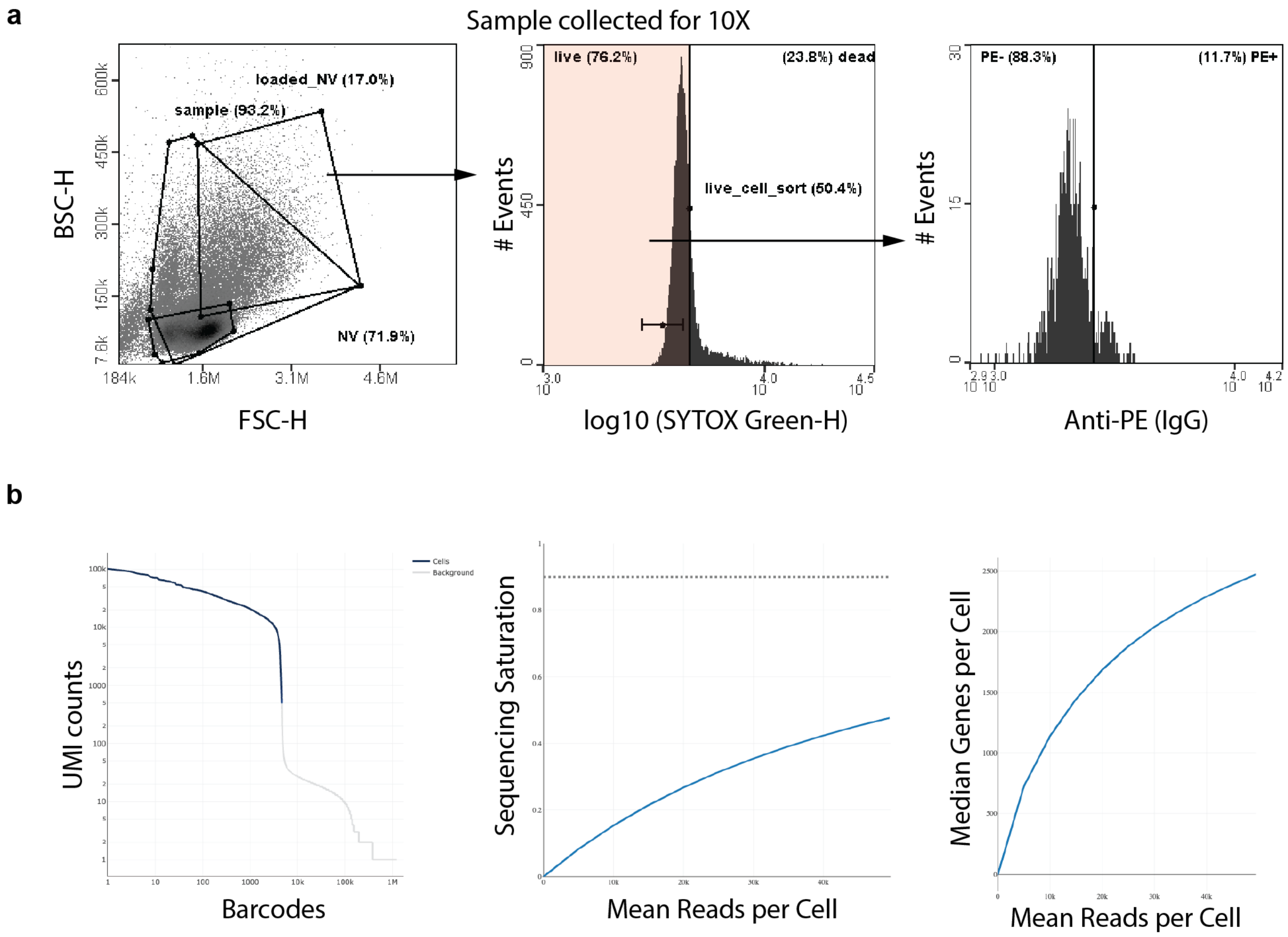


**Figure S8. Sorting live-cell-containing nanovials for SEC-seq and Cell Ranger QC**

**(a)** Sorting strategy for live-cell-containing nanovials using the Nanocellect WOLF prior to loading into the 10X Chromium Next GEM chip. All live cells that were loaded on nanovials were collected independent of secretion amount. Analysis of the sorted population showed a distribution of secretors with ~12% of the population above the marked threshold. **(b)** 10X Cell Ranger QC summary. Barcode rank plot represents the UMI count separation between cells and background (left). Sequencing saturation plot shows the sequencing depth is ~50% by using a NextSeq 1000/2000 kit (middle). Median genes per cell plot shows the full sequencing depth is about 2500 median genes per cell.

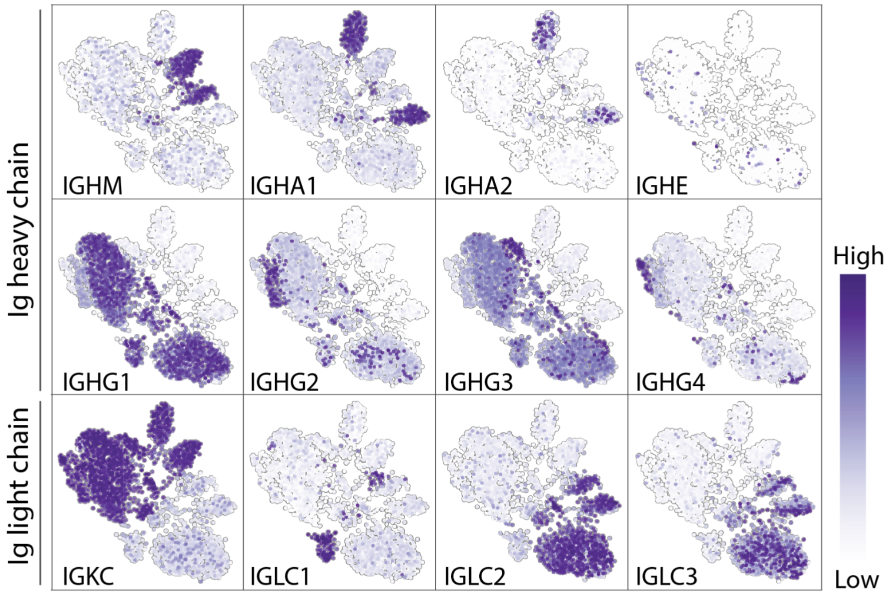


**Figure S9. UMAP of immunoglobulins.**
Transcript levels of immunoglobulin genes projected on the UMAP plot from Figure 4b. The top row represents non-IgG immunoglobulin classes, the middle row represents IgG subclasses, and the bottom row represents immunoglobulin light chain.

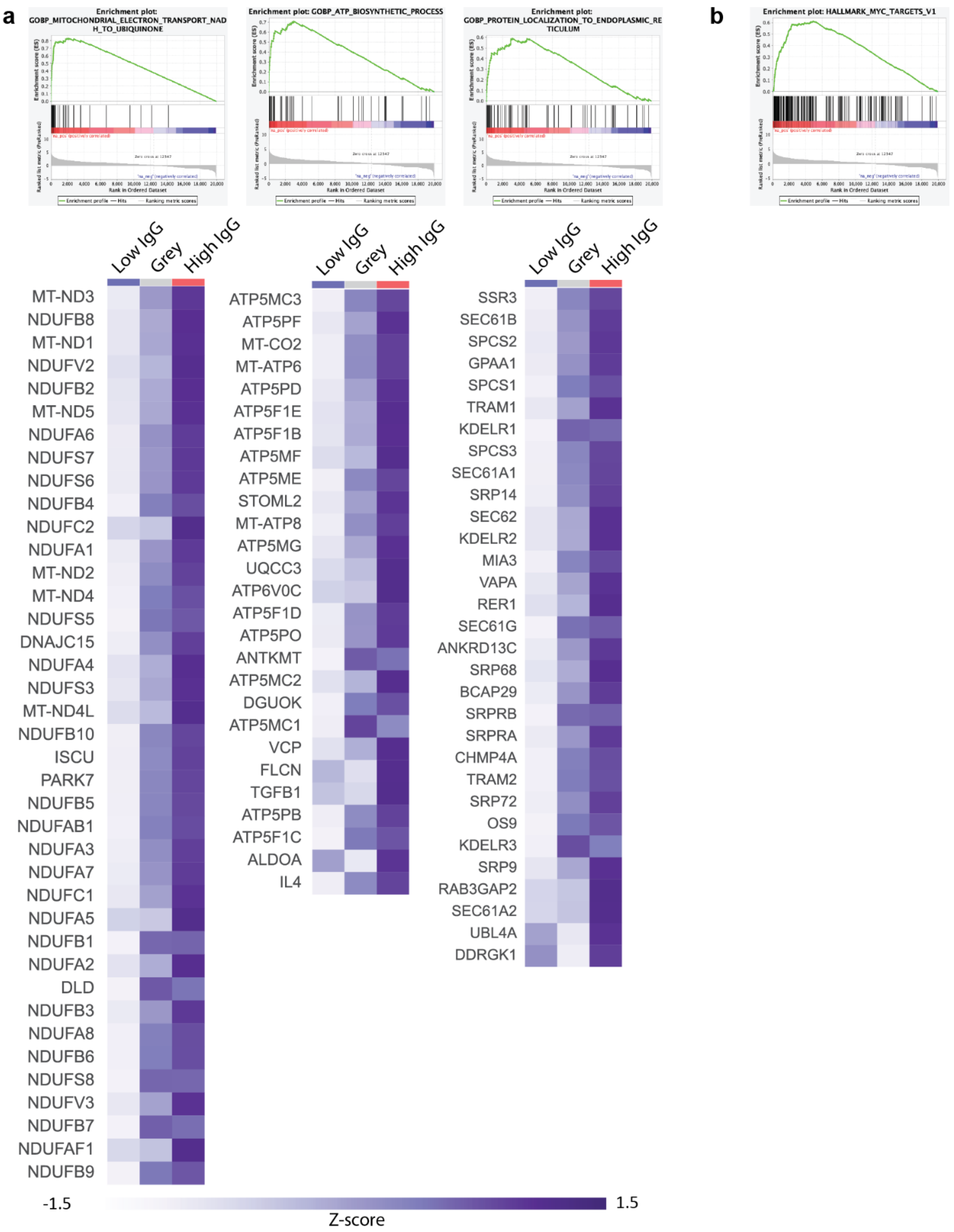
a
Enrichment plot: GOBP_MITOCHONDRIAL_ELECTRON_TRANSPORT_NADH_TO_UBIQUINONE
Enrichment plot: GOBP_ATP_BIOSYNTHETIC_PROCESS
Enrichment plot: GOBP_PROTEIN_LOCALIZATION_TO_ENDOPLASMIC_RETICULUM
b
Enrichment plot: HALLMARK_MYC_TARGETS_V1
Low IgG
Grey
High IgG
MT-ND3
NDUFB8
MT-ND1
NDUFV2
NDUFB2
MT-ND5
NDUFA6
NDUFS7
NDUFS6
NDUFB4
NDUFC2
NDUFA1
MT-ND2
MT-ND4
NDUFS5
DNAJC15
NDUFA4
NDUFS3
MT-ND4L
NDUFB10
ISCU
PARK7
NDUFB5
NDUFAB1
NDUFA3
NDUFA7
NDUFC1
NDUFA5
NDUFB1
NDUFA2
DLD
NDUFB3
NDUFA8
NDUFB6
NDUFS8
NDUFV3
NDUFB7
NDUFAF1
NDUFB9
ATP5MC3
ATP5PF
MT-CO2
MT-ATP6
ATP5PD
ATP5F1E
ATP5F1B
ATP5MF
ATP5ME
STOML2
MT-ATP8
ATP5MG
UQCC3
ATP6V0C
ATP5F1D
ATP5PO
ANTKMT
ATP5MC2
DGUOK
ATP5MC1
VCP
FLCN
TGFB1
ATP5PB
ATP5F1C
ALDOA
IL4
SSR3
SEC61B
SPCS2
GPAA1
SPCS1
TRAM1
KDELR1
SPCS3
SEC61A1
SRP14
SEC62
KDELR2
MIA3
VAPA
RER1
SEC61G
ANKRD13C
SRP68
BCAP29
SRPRB
SRPRA
CHMP4A
TRAM2
SRP72
OS9
KDELR3
SRP9
RAB3GAP2
SEC61A2
UBL4A
DDRGK1
-1.5
1.5
Z-score

**Figure S10. GSEA plot and gene list in core enrichment.**
**(a)** Representative GSEA plots and gene lists in core enrichment in GO biological process: Mitochondrial electron transport NADH to ubiquinone (left), ATP biosynthesis process (middle) and Protein localization to endoplasmic reticulum (right). **(b)** GSEA plot of Hallmark MYC-targets V1 that have the expanded gene list in Figure 6.
